# Mechanism and energetics of JDP induced Hsp70’s conformational transition towards catalytically active state

**DOI:** 10.1101/2025.05.22.655504

**Authors:** Michał Olewniczak, Marcin Pitek, Jacek Czub, Jarosław Marszałek, Łukasz Nierzwicki, Bartlomiej Tomiczek

## Abstract

Hsp70 chaperones are crucial for maintaining protein homeostasis by regulating the stability and conformational states of client polypeptides through cycles of their binding and release. These cycles require conformational transitions of Hsp70 driven by ATP binding and hydrolysis. The ATPase activity of Hsp70 is controlled by J-domain protein (JDP) cochaperones, which allosterically stimulate ATP hydrolysis via interactions between their J-domains and Hsp70. The J-domain binds at the interface between the nucleotide (NBD) and substrate (SBD) binding domains of ATP bound Hsp70. Although, it was established that the JD interaction involves residues of helices II and III, and the interhelical loop critical for ATPase stimulation, the mechanism by which the allosteric signal induced by J-domain binding is transmitted to the distal nucleotide-binding pocket of Hsp70 remains unclear, as do the conformational changes leading to the ATP hydrolysis. Here, we addressed these questions by means of all-atom free energy simulations and dynamic network analysis, starting from the crystal structures of ATP-bound Hsp70 DnaK alone and in complex with the J-domain of DnaJ. We demonstrated that the presence of the J-domain results in the rearrangement of the nucleotide-binding pocket into a hydrolysis competent state, characterized by close contact between universally conserved T199 of NBD and γ-phosphate of ATP. With network analysis we revealed that the allosteric signal for this rearrangement is transmitted along the β-strand containing T199. Finally, we provide rationale for the signal transmission, where steric repulsion between the J-domain’s helix III and SBD induces a push of the T199 containing β-strand. Overall, our study provides mechanistic insights into allosteric signal transmission within Hsp70, bridging the gap between J-domain binding and ATPase stimulation.

**TOC Graphic:** 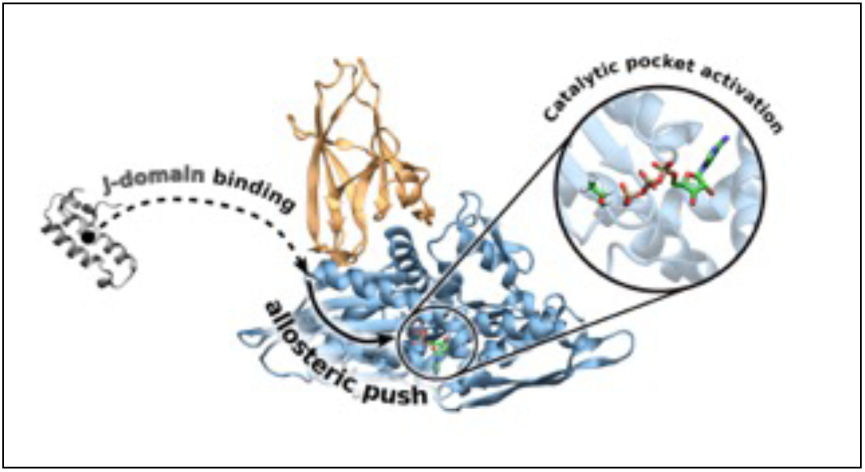

## Introduction

Cells are frequently exposed to environmental stressors, such as temperature, oxidative stress, toxic substances or radiation, which can disrupt protein homeostasis and cause protein aggregation, leading to cell death. Hsp70 chaperones together with J-domain protein (JDP) cochaperones play a critical role in controlling conformational state of other proteins. Hsp70 machinery promotes protein folding into functional native states through ATP-dependent cycles of client protein binding and release^1^. The client binding cycle of Hsp70s is strictly controlled by interactions with the J-domain (JD) of their JDP cochaperone partners, which stimulate ATP hydrolysis, facilitating the transition from an ATP-bound state to an ADP-bound state.^2^ In the ATP-bound state, the nucleotide binding domain (NBD) of Hsp70, which binds and hydrolyses ATP, is tightly docked to the interdomain linker and the substrate binding domain (SBD)(Figure 1). JDP binds client proteins and recruits Hsp70 in the ATP-bound state.^3^ Binding of both the client and JD to Hsp70 stimulates ATP hydrolysis by inducing the allosteric transition from the restraining state (R), where ATP hydrolysis is restricted, to the stimulating state (S), where hydrolysis occurs rapidly. Such transition was previously observed for the client, and although it is known that both JD and client stimulate Hsp70 ATPase activity in a synergistic manner, the conformational transition upon JD binding is not fully characterized and lacks mechanistic explanation.^4^ After hydrolysis, the SBD undocks from the NBD and the α-helical lid (SBDα) starts to interact with ϕ3-sandwich domain of SBD(SBDϕ3), stabilizing the Hsp70-client interaction.^5^ Finally, after JD dissociation from Hsp70, the ADP-to-ATP exchange is facilitated by another class of co-chaperones termed Nucleotide Exchange Factors (NEFs), and ATP rebinding triggers conformational changes back to the Hsp70 ATP-bound state. ^6–8^

**Figure 1.**
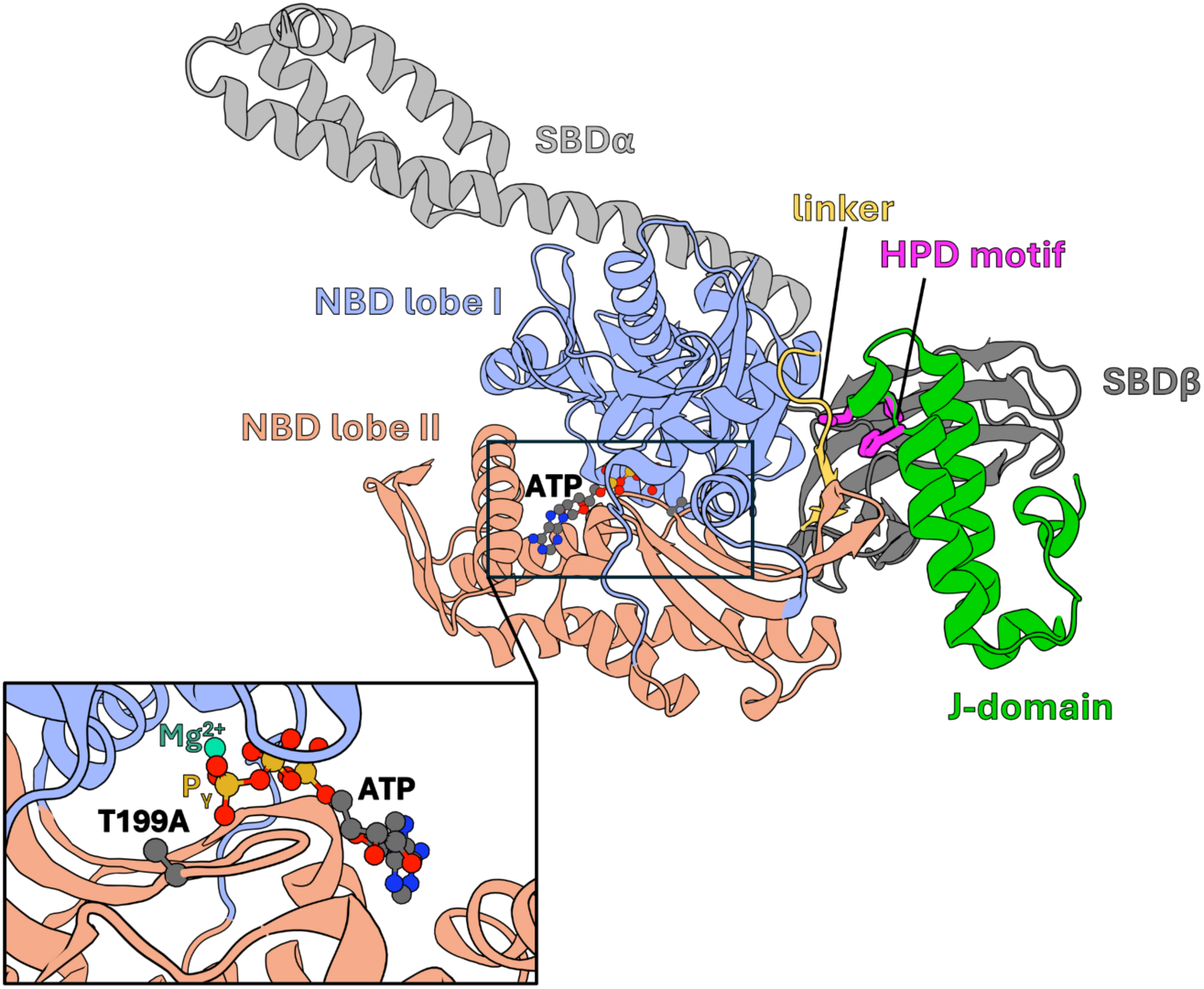
Structure of DnaK – DnaJ J-domain complex (PDBID: 5NRO) in cartoon representation (NBD lobe I – light blue, NBD lobe II – light red, interdomain linker – yellow, SBDβ – dark gray, SBDα – light gray, J-domain – green). J-domain’s HPD motif in licorice representation – magenta. ATP in CPK representation colored according to the elements (carbon – dark grey, nitrogen – blue, oxygen – red, phosphorus – orange). The insert shows the position of T199A substitution in relation to the ATP and Mg^2+^(teal).

The mechanism by which the JD allosterically communicates with the residues of the ATP binding site remains puzzling despite extensive studies.^9–11^ Substituting threonine 199 with alanine (T199A) in the nucleotide-binding site impairs the ATPase activity of Hsp70,^10,12,13^ a residue which typically contributes to stabilizing the metaphosphate intermediate in many biomolecular motors.^14,15^ While a crystal structure with the T199A mutation (PDB: 5NRO) revealed a network of contacts connecting the JD to the catalytic center of the NBD, crystallographic data offer only limited insights into the mechanistic importance of T199 in ATP hydrolysis, and the associated conformational changes remain unclear^5^. Previously, we demonstrated that JD binding leads to the displacement of R167 in the NBD by the highly conserved HPD motif of the JD, triggering the conformational transition of Hsp70.^10,12^ The JD is sufficient to provide a signal for the ATP hydrolysis and acts synergistically with client proteins to stimulate Hsp70’s ATPase activity.^13^ However, studies on the *S. cerevisiae* Hsp70/JDP (Ssq1/Hsc20) system showed that alanine substitutions of the HPD motif are not sufficient to fully abolish the JD-induced ATPase activity^14^ challenging the current understanding of the allosteric stimulation in Hsp70s by JDPs.

To shed light on the molecular mechanism by which JD transmits the allosteric signal for ATP hydrolysis within the NBD of Hsp70, we reverted T199A substitution in the crystal structure of Hsp70 and Hsp70-JD complex (DnaK/DnaJ) from *E.coli* and performed all-atom free-energy simulations. We characterized the conformational changes in Hsp70 upon JD binding that led to a T199 displacement towards γ-phosphate of the ATP. Our results also highlighted the role of the HPD motif in ensuring the proper orientation of JD with respect to Hsp70. Furthermore, using dynamic network analysis, we showed that communication between T199 and the JD is mediated by the lobe II of the NBD and identified allosteric pathways through which signal is propagated. Finally, analysis of the truncated variants of JD allowed us to explain the role of helix III in initiating conformational transition of Hsp70.

## Results and discussion

### JD induces conformational changes in Hsp70

J-domain (JD) interacts with ATP-bound Hsp70 and is thought to allosterically promote conformational transition of NBD to the stimulating state competent for ATP hydrolysis.^4,15,16^ Therefore, to characterize the essential motions of Hsp70 upon JD binding, we performed Principal Component Analysis (PCA) of the Hsp70 (DnaK) alone and in complex with JD (DnaK-JD). For that purpose, both structural ensembles were obtained using all-atom molecular dynamics (MD) simulations with restored T199 residue. We found that the first principal component, which describes the most pronounced collective motion of NBD (28% of the total conformational variability; Figure S2), was sufficient to distinguish the NBD states of Hsp70 alone and in complex with the JD (Figure S1). Notably, the values of the first principal component were highly correlated with the torsion angle between the NBD lobes (R^2^=0.95), indicating that upon JD binding, the angle between NBD lobes decreases by ∼7 ° compared to Hsp70 alone resulting in a more compact state of the NBD (Figure 2A,B).

**Figure 2.**
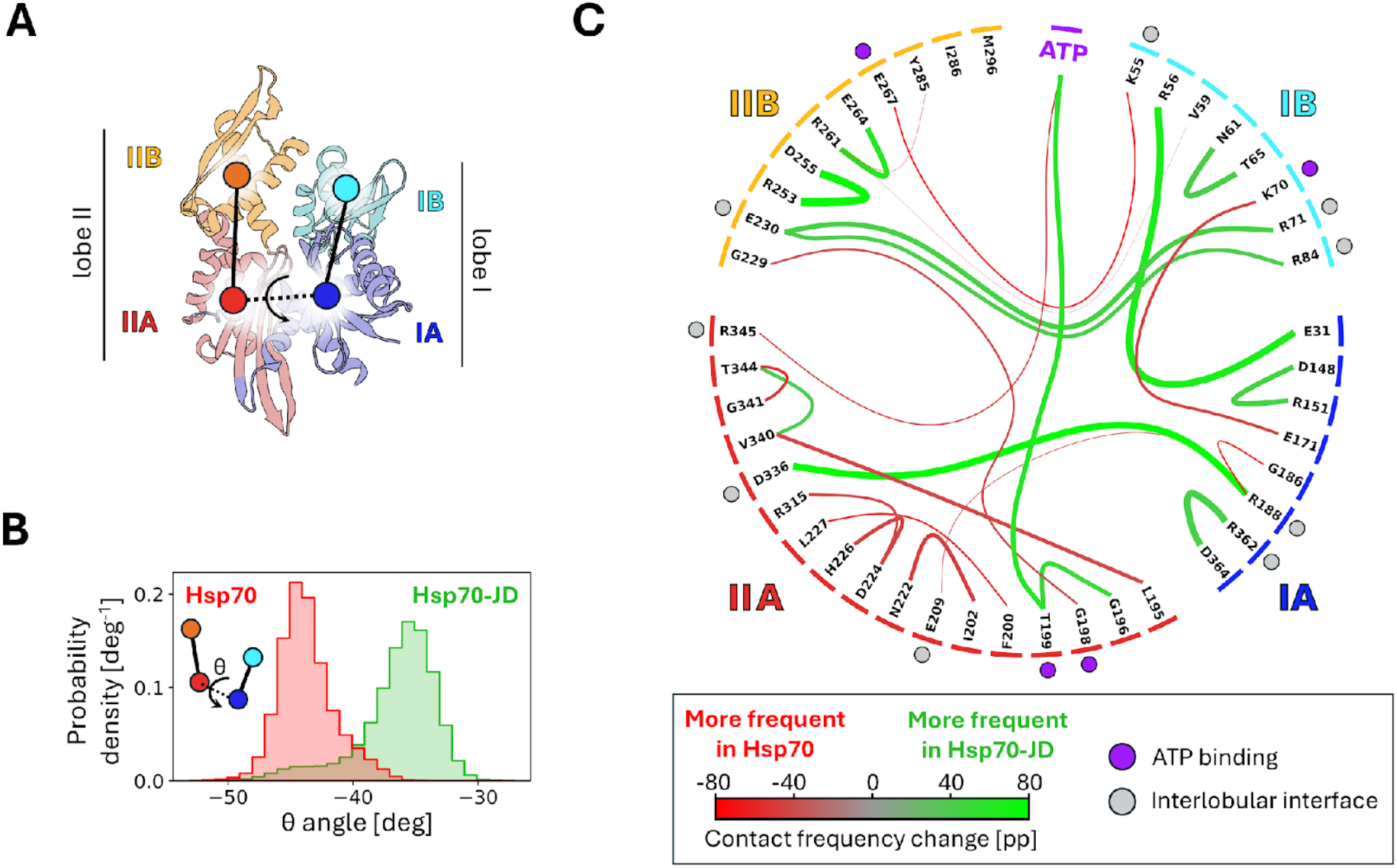
(A) Structure of the Hsp70’s NBD (DnaK) colored by the subdomains (PDBID: 5NRO). The subdomain IA of NBD is depicted in blue, IB– in cyan, IIA– in red and IIB– in orange. Dots indicate subdomain centers of mass, used to define torsion angle describing subdomain dynamics. (B) Probability distribution of the torsion angle between the NBD subdomains during MD simulations of the Hsp70 (red) and Hsp70-JD complex (green). (C) Chord diagram showing changes in NBD residues contact frequency associated with Hsp70-JD complex. Line widths are proportional to the contact frequency in Hsp70-JD complex and are colored based on the contact frequency difference with respect to Hsp70. Only contacts displaying frequency difference of 40 percentage points or greater are shown. Dots positioned next to the NBD residues indicate ATP-binding residues (violet) or residues positioned at the interlobular interface (gray) in the Hsp70-JD complex. Colored bars along the circle’s rim indicate NBD subdomain.

To further characterize the conformational changes of Hsp70 resulting from the JD binding, we compared the frequency of inter-residue interactions, including hydrogen bonds, ionic interactions, *π*-stacking, *π*-cation interactions and hydrophobic interactions in Hsp70 alone and in the Hsp70-JD complex, focusing on the NBD (Figure 2C). Notably, the most significant differences in contact frequency were observed in: (i) the JD binding site of the NBD; (ii) the ATP-binding site of the NBD and (iii) lobe IIA (see also Figure S3-S4). Strikingly, we observed that in the Hsp70-JD complex the interaction between the hydroxyl group of T199, a residue critical for Hsp70’s ATPase activity^17^ and ATP is present, while absent in the Hsp70 alone (Figure S5). In the Hsp70-JD complex, T199 contacts γ-phosphate of ATP, similarly as in many ATPases,^18,19^ where it plays a role in stabilizing the metaphosphate intermediate during ATP hydrolysis. Additionally, our simulations of the Hsp70-JD complex captured other local changes in the nucleotide binding site. Specifically, the hydroxyl group of residue Y145 is displaced from the SBDβ interface to the ATPase catalytic site, possibly stabilizing T199 positioning close to the γ phosphate of ATP (Figure S6 shows probability distribution of the distance between tyrosine hydroxyl (hydrogen bond donor) and the hydroxyl of T199 (hydrogen bond acceptor)).

Taken together, our MD simulations of the ATP bound Hsp70-JD complex with restored T199 revealed the allosteric effect of the JD, which involves conformational transitions within the NBD. These transitions consist of rotation of the NBD lobes and reorganization of the NBD contact network, including the displacement of T199 towards γ−phosphate of ATP and Y145 towards T199. This observation is in good agreement with recent studies characterizing the client-induced transition of Hsp70 between the restraining (R) and stimulating (S) states, which involves similar rotation of the NBD lobes and displacement of Y145 residue toward the γ-phosphate of ATP in absence of T199.^4^ These results suggest that Y145 displacement is a common step in client- and JD-induced stimulation of Hsp70, possibly accounting for their known synergistic effect. At the same time, a number of residues previously found to play a role in the allosteric communication between the NBD and SBD (K70, D148, R151, E171) ^17,20,21^ exhibit substantial changes in their interaction patterns in the Hsp70-JD complex compared to Hsp70 alone (Figure S4).

### JD facilitates the positioning of T199 next to the γ-phosphate of ATP

We examined the structural determinants of T199 positioning within the catalytic site of Hsp70. Specifically, we ask if JD binding favors the displacement of T199 towards γ-phosphate of ATP. To quantitatively address this question, we characterized the thermodynamics of the T199 conformational change by means of enhanced sampling methods (Figure 3). We determined the free-energy profiles for T199 movement towards ATP γ-phosphate for both Hsp70-ATP alone and in complex with the JD. The free energy changes with the distance between T199 hydroxyl oxygen and the γ-phosphate (T199-γP) of ATP were sampled either in the presence or absence of the JD. We used a well-established umbrella sampling method to determine the free-energy profiles and ran the simulations extensively until convergence. Comparison of the two resulting energy profiles reveals that T199 positioning strongly depends on the presence of the JD. The free energy profile for Hsp70 alone shows a well-defined minimum corresponding to the T199 hydroxyl group being displaced from the γ-phosphate by ∼0.8 nm (Thr-far state). Formation of the T199 - γ-phosphate contact in that state is associated with an energy cost of ∼6 kcal/mol. We thus conclude that in the absence of JD, T199 sidechain is displaced from the γ-phosphate and cannot participate in stabilization of the ATP-to-ADP transition state. In contrast, in the Hsp70-JD complex, the free energy profile shows a pronounced global minimum at ∼0.35 nm corresponding to the Thr-close state. Formation of the T199 - γ-phosphate contact in Hsp70-JD complex is favorable and associated with an energy gain of ∼3 kcal/mol.

**Figure 3.**
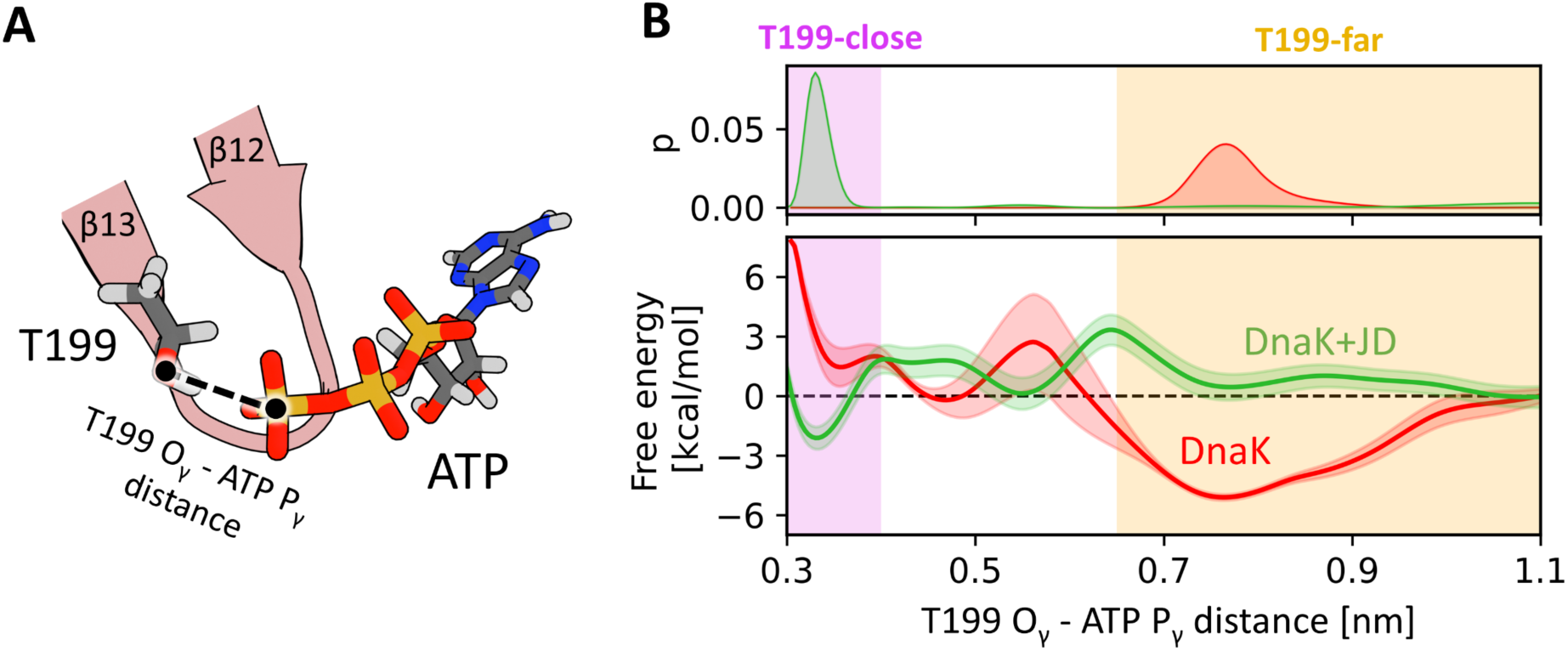
(A) Schematic representation of the reaction coordinate used in the umbrella sampling simulations. The atoms shown in licorice representation are coloured according to the elements – carbon in dark grey, oxygen in red, hydrogen in light grey, phosphorus in orange, nitrogen in blue. (B) Probability distributions (top) and free energy profiles (bottom) for the distance between hydroxyl oxygen of T199 and the phosphorus of ATP gamma phosphate in the absence (red) and presence (green) of the JD. The purple shade indicates the Thr-close state (distance < 0.4 nm), whereas orange shade indicates the Thr-far state (distance > 0.65 nm).

This result indicates that the Thr-close state, is adopted most frequently in the presence of JD, in contrast to the Hsp70 alone where the Thr-far state is favorable, as indicated by the probability distribution along the T199 - γ-phosphate distance (Figure 3B). Without JD, Thr-close state is not easily accessible since it is 6 kcal/mol less favorable than the Thr-far state. Overall, our results support the hypothesis that JD binding induces the rearrangement of the ATP binding site via induced fit mechanism, enabling T199 interaction with the γ − phosphate of ATP and accounting for the ATPase stimulation.

### Allosteric signaling between JD and catalytic site of Hsp70

To better understand how the JD interaction at the surface of Hsp70 induces conformational changes within the catalytic site located on the other side of NBD ,^10,12^ we first tested the essentiality of residues within HPD motif of the JD for the allosteric signaling. The role of HPD motif in stimulation of ATP hydrolysis is well established^10,22,23^, although the exact functions of individual residues within the motif are not fully resolved. To address this, we generated single alanine substitution variants within the HPD (APD, HAD, HPA), and performed unbiased MD simulations. In the Hsp70-JD(H33A) complex, the dissociation of JD was not observed, and the equilibrium state closely resembled that of the native Hsp70-JD complex. Additionally, the computed free energy profile for the binding of H33A JD to Hsp70 was almost identical to that of the native JD (Figure S14; left). We then investigated whether the H33A mutant also causes the same displacement of T199 in the catalytic site of Hsp70 as the native JD. As before, using umbrella sampling, we determined how the free energy changes with the distance between T199 hydroxyl oxygen and the terminal phosphorus atom of ATP. Remarkably, with the H33A substitution, the Thr-far state is 3 kcal/mol more stable than the supposedly catalytically active Thr-close state (Figure S14; right). This result shows that, although H33A mutation does not lower the binding affinity of the JD for Hsp70, it promotes the Thr-far state and thus prevents the conformational transition of Hsp70 to a stimulating state.

Next, we analyzed the effects of D35A substitution. Our observations indicate that JD partially dissociates from the NBD binding site leaving only its helix II in direct contact with the NBD. In the Hsp70-JD(D35A) complex, both helix III and loop between helix II and III (HPD loop) have a higher probability of being at a greater distance from Hsp70 binding site, compared to the native Hsp70-JD complex (Figures S13, S14). Further, we calculated free energy profiles along the distance between the Hsp70 binding site of JD and the JD using umbrella sampling simulations. The obtained profiles (Figure S14; left) clearly indicate that substituting D35 with alanine results in JD binding that is by ∼2 kcal/mol less favorable and causes a relaxation in the complex structure, as indicated by the free energy ‘minimum located ∼0.2 nm further along the Hsp70-JD distance. We also analyzed the effects of the D35A substitution on the destabilization of the SBD*β*-NBD interface. Consistently with our previous studies, JD binding to Hsp70 causes partial displacement of SBD*β* from NBD.^12^ Yet, additional umbrella sampling simulations along the SBD*β*–NBD distance for the Hsp70-JD(D35A) complex revealed that the D35A variant of JD does not induce this displacement (Figure S17). These results highlight the critical role of the salt bridge between D35 and R167^12^ in establishing the conformation proper for ATP hydrolysis of the Hsp70-JD complex. Therefore, although the effect of D35A and H33A mutations on JD binding is markedly different, they both abolish the conformational transition of Hsp70 to the stimulating state, but through different mechanisms.

Since H33 and D35 in the HPD motif directly interact with the specific amino acids of NBD (L391 and R167, respectively), we hypothesized that local disruption of alpha helix in JD by proline 34, commonly known to act as a secondary structure breaker^24^, enables H33 and D35 to adopt orientation favoring interaction with Hsp70. To test this hypothesis, we performed MD simulations of the free JD and its P34A variant. As expected, we observed that the P34A substitution extends helix II up to the HPD motif. Moreover, in the native JD, a dihedral angle describing the relative orientation of H33 and D35 adapts values close to 0 **°**, indicating that both residues are facing the same direction (Figure S11). The JD(P34A) variant adopts a broad value distribution of the dihedral angle between H33 and D35 residues, contrary to a narrow one observed in the Hsp70-JD bound state.

To elucidate allosteric signal transmission routes transmitted from the HPD motif in the JD to the T199 in the Hsp70 catalytic site, we constructed a dynamic network model, representing a protein as a network of C*α* atoms (nodes) connected by edges, with lengths determined by the strength of motion correlations between residues. Our multi-µs MD trajectory analysis revealed that all identified allosteric pathways follow the same pattern: the signal originating from D35 is mediated through the Hsp70 interdomain linker and NBD *β*-strands 13 and 14 (residues F200– T215), finally reaching T199 (Figure 4, S7). Notably, 7 out of 14 residues participating in these pathways were previously suggested to contribute to the allosteric communication within Hsp70 in mutational and structural studies.^20,23,25–27^ In contrast to other studies, none of the pathways were mediated by R167, which was previously proposed to play an important role in allosteric communication towards ATP hydrolysis in Hsp70.^10^ Because in Hsp70-JD(D35A) complex was relaxed, we determined allosteric pathways for previously described Hsp70 (R167A) - JD (D35N) compensatory mutant. Notably, all identified pathways followed similar pattern as the wild type complex (Figure S21). Additionally, we performed free energy calculations between Thr-far and close states for the Hsp70 (R167A)-JD (D35N) variant. Our energy profile shows the flattening of the free energy landscape between Thr-far and close states, making the latter one more accessible than in Hsp70 alone (Figure S20, S21). This explains why simultaneous replacement of R167 and D35 by alanine and asparagine, respectively, is known to partially restore the Hsp70 function.^28,29^

**Figure 4.**
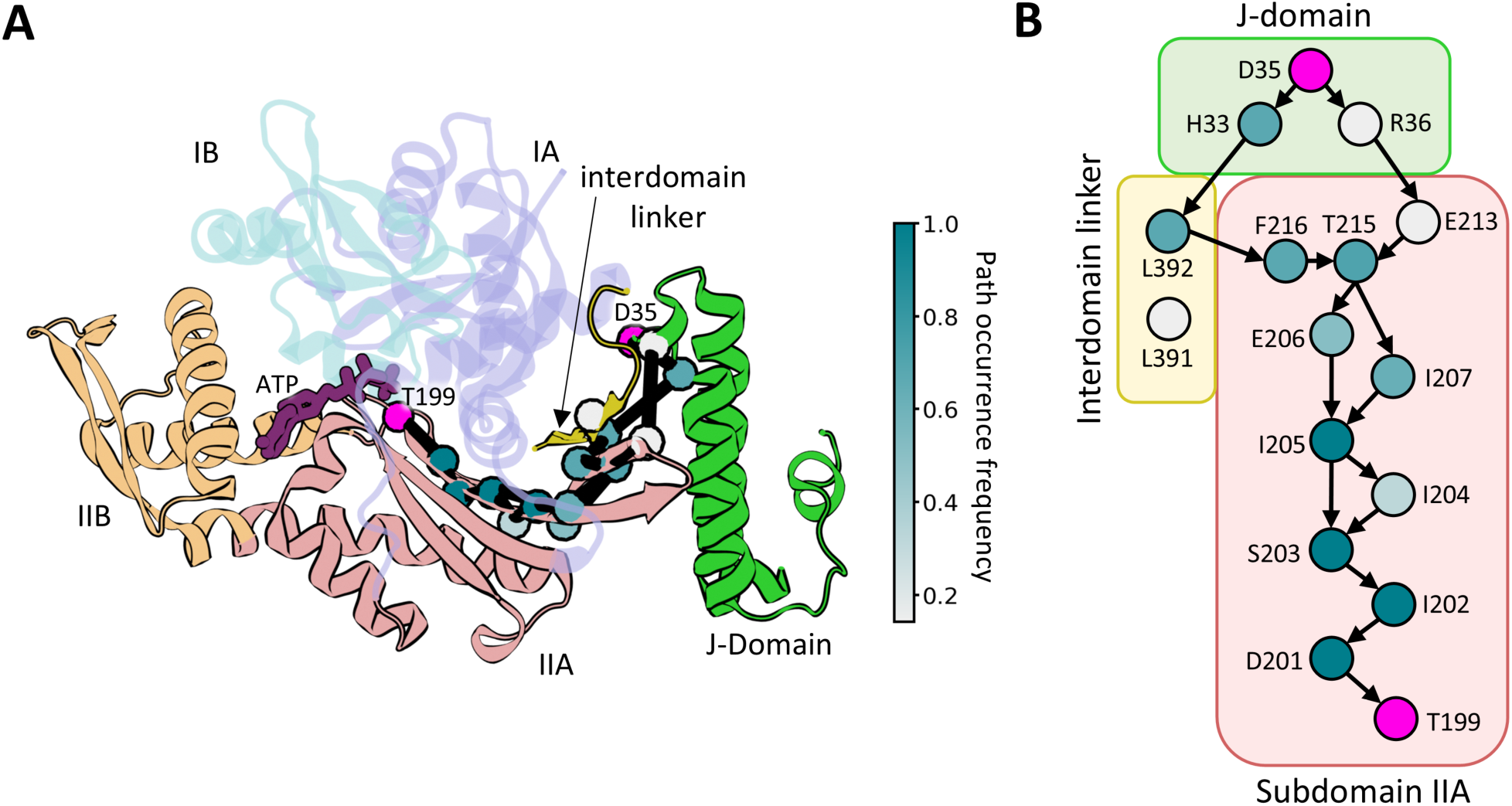
Allosteric signaling between JD and Hsp70. (A) Set of 5 optimal pathways between the D35 of the JD HPD motif and the T199 of Hsp70 (magenta spheres) calculated for 5 Hsp70-JD MD trajectories and projected onto the Hsp70-JD complex. C_α_ atoms of residues present in the set of optimal and suboptimal pathways shown as spheres and colored according to the path occurrence frequency (for more details see Figure S7). (B) Graph depicting connectivity of pathway forming residues. Arrows represent connections present in the set of 5 optimal pathways, residue coloring as in (A).

To further evaluate the closeness of the JD to the T199 in the correlation space, we employed the community network analysis,^30,31^ which simplifies the dynamic network by clustering the strongly connected nodes into communities. The obtained community structure roughly reflects the division of the NBD into subdomains (Figure S8– S9). The T199 residue was assigned to a zeroth community, which is directly connected to the first one containing the JD. This helps to explain the observed routing of paths between D35 of the JD and T199 as minimization of community crossings by the path should be beneficial to its length in the correlation space. Additionally, three leucines in the conserved VLLL motif within the interdomain linker were assigned to the same community as JD, highlighting the tight contact between these regions. We then quantified the strength of the connections between the communities using the betweenness centrality measure (BC; see Methods for more details) and analyzed pivotal interactions for allosteric signal transmission based on the edges with the highest BC. Both subdomain IA and IIA formed strong connections with the JD. Specifically, residues D35, K26, Y25, H33 and F47 of JD interact with R167, Q217, L391 and L392 residues of NBD. Notably, D35 and H33 of the HPD make direct contact with R167 of NBD and L391 of the interdomain linker, respectively (Figure S10). H33– directly interacts with L391 of the interdomain linker and participates in transmitting of the allosteric signal to the catalytic site of Hsp70 through L392 or L391 (Figures 4, S10). ^10^

### JD-induced allosteric signaling is associated with destabilization of Hsp70’s β-strand 14

To further characterize conformational changes, that are directly responsible for the displacement of T199 towards ATP γ-phosphate (Thr-close) facilitating ATP hydrolysis, we performed a series of conventional MD simulations of Hsp70-JD complex. We observed multiple events where Thr-close state in the catalytic site of Hsp70 was associated with the disruption of *β*-strand 13 (which includes T199) - *β*-strand 14 interaction (residues 198-206 and 219-227; Figure S18A) and disruption of *β*-strand 14 conformation. These two *β*-strands are part of *β*-sheet together with *β*-strands 12, 17 and the linker. Strand 14 which is part of “switch segment” together with G-loop connecting *β*-strand 12 and 13 were previously identified as main conformational differences between the restraining (R) and stimulating (S) states of Hsp70.^4^ When we compared fraction of residues adopting *β*-sheet conformation in the structural ensemble of isolated Hsp70 and the Hsp70-JD complex, the probability distribution indicated that interaction between these two *β*-strands is more frequently disrupted in the Hsp70-JD complex than in Hsp70 alone (Figure S18B).

To examine how this conformational change affects the interaction between T199 and ATP, we performed a set of free energy simulations along the T199–ATP distance, while considering two different conformations of the G198-E206/L219-L227 *β*-region. First, we enforced disrupted conformation of the fragments of *β*-strands 13 and 14 in the Hsp70 system to mimic the effect of JD (see Methods for more details). The obtained profile (Figure S16; left) is highly similar to that of isolated Hsp70, suggesting that *β*-strands disruption alone does not directly lead to the Thr-close state. Second, we enforced undisrupted conformation of the *β*-strand in the Hsp70-JD complex. The corresponding profile along the T199–ATP distance shows a clear preference for adopting the Thr-far state (Figure S16; right). Because tight JD binding causes the displacement of the NBD and SBD*β* subdomains, it could explain why forced disruption of the β-strand in the absence of JD was insufficient to favor the Thr-close conformation. Conversely, our results for the D35A JD variant already indicated that displacement of the NBD lobes does not occur when JD is loosely bound to Hsp70, i.e. when the helix III of JD dissociates from the Hsp70 surface. We therefore hypothesized that in the Hsp70-JD complex, the helix III of tightly bound JD induces conformational changes in Hsp70 and leads to disengagement of NBD from the SBD. In this scenario, the contact force exerted by the JD could be the cause for both displacement of SBD*β* from NBD and the β-sheet disruption.

To support this hypothesis, we constructed an Hsp70 complex with JD lacking helix III (residues 41-65) to evaluate how this change would affect the energetics of a) SBD*β* and NBD dissociation and b) the close interaction of T199 with ATP. The resulting free energy profiles for the JD ΔH3-4 variant (Figure 5) are similar to those for Hsp70 alone and indicate that the lack of helix III of JD prevents the SBD*β*-NBD displacement thus stabilizing the Thr-far conformation, supporting our hypothesis that the contact force exerted by helix III is necessary for transition towards Thr-close state.

**Figure 5.**
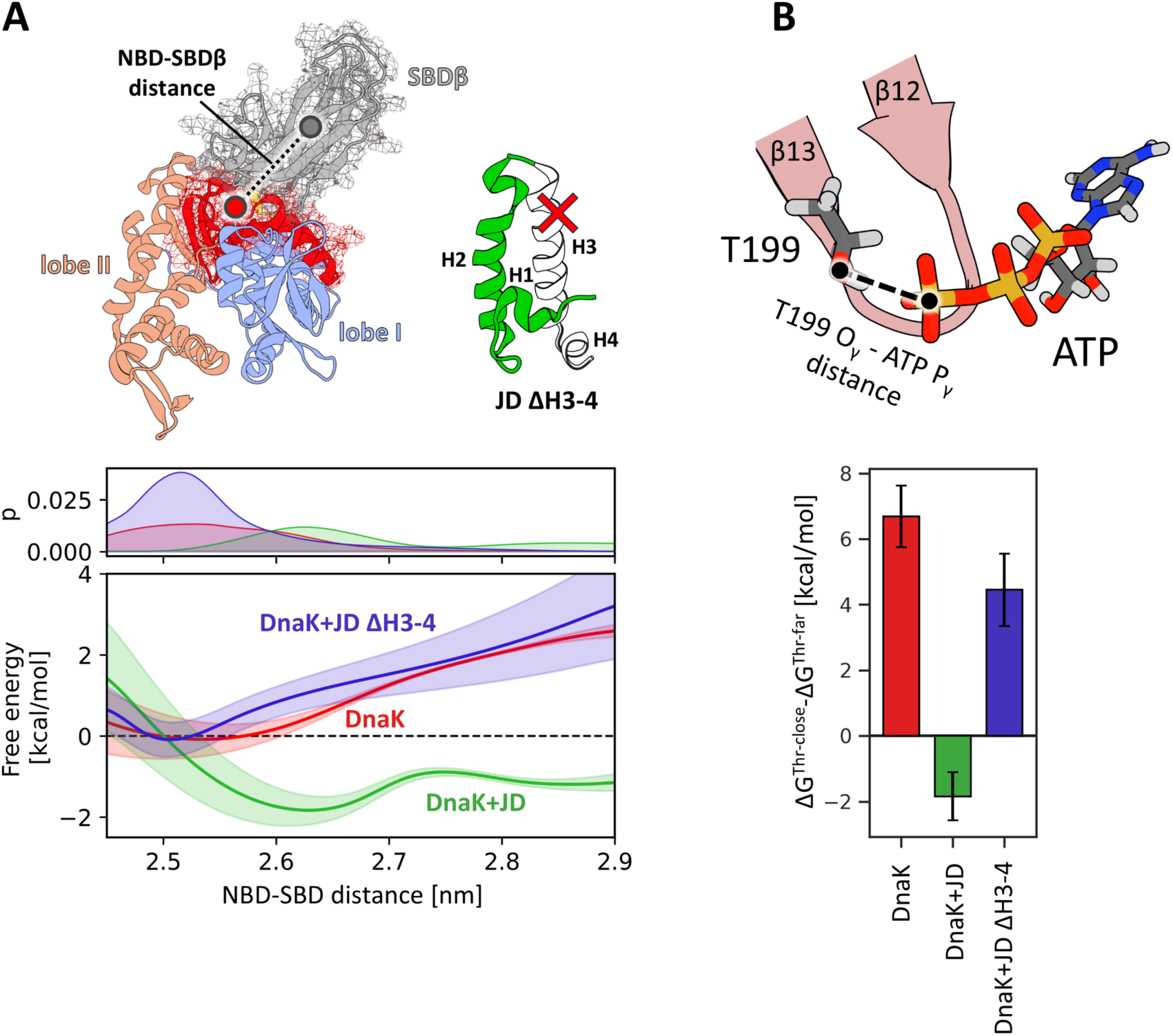
Role of the helix III of JD. (A) Free energy profiles for the dissociation of the NBD and SBD*β*. The profile for the Hsp70 without JD is depicted in red, for the complex of Hsp70 and WT JD– in green, whereas for the complex of Hsp70 with the JD lacking helix III and IV– in blue. The definition of the distance between centers of mass of the SBD*β* and the part of NBD interfacing SBD*β* is shown above. (B) Differences of the free energy associated with the transition from the Thr-far to Thr-close state for the same systems as in 4A (for the free energy profiles see Figure S15).

Taken together, our results support a model, in which helix III of JD acts like a wedge, facilitating the formation of the optimal interface between the SBDβ and NBD and this interaction is required for maintaining Thr-close state (Figure 6). The resulting strain on the NBD causes disruption of the *β*-strands 13 and 14 (G198-E206/L219-L227 beta-sheet), which stimulates the ATPase activity of Hsp70 by favoring the close contact of T199 with ATP γ-phosphate and stabilizing the transition state during hydrolysis. Importantly, as illustrated by the behavior of the D35A JD variant, to perform its role, helix III requires the tight binding of JD to Hsp70 ensured by the HPD aspartate.

**Figure 6.**
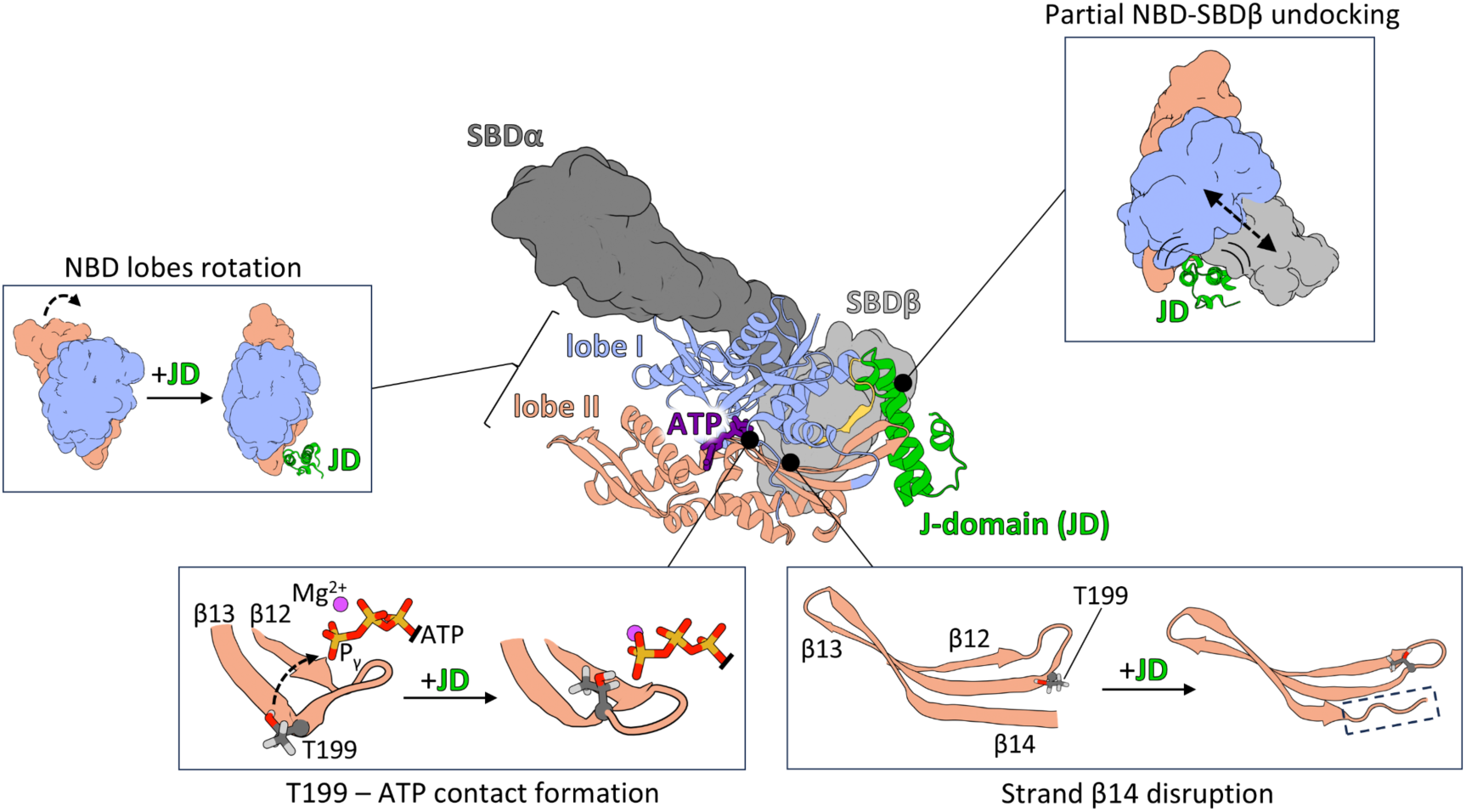
Schematic representation of the mode of action of the JD in the stimulation of ATPase activity. By tight binding to Hsp70, JD exerts both local and global conformational changes within Hsp70. Globally JD promotes partial NBD-SBDβ undocking, which diminish the SBDβ inhibitory effect on NBD lobes rotation reducing the torsion angle between them. Locally the JD binding initiates propagation of allosteric signal through β-sheet in subdomain IIA, which results in contact formation between T199 and gamma phosphate of ATP and disruption of β-sheet’s strand β14. Altogether observed rearrangements prime the Hsp70’s NBD for the ATP hydrolysis. The SBD*β* is depicted in light grey, SBDα – in dark grey, NBD lobes I and II – in light blue and salmon respectively and the JD– in green. The atoms shown in licorice representation are colored according to the elements – carbon in dark grey, oxygen in red, hydrogen in light grey, phosphorus in orange and magnesium in magenta.

## Conclusions

In this study, we investigated the mechanism of Hsp70s stimulation by their co-chaperone J-domain proteins (JDPs), using Hsp70/JD pair from E.coli, DnaK and DnaJ, as a model. Our extensive unbiased simulations showed that the binding of JD results in the rearrangement of the NBD lobes in Hsp70. Specifically, the presence of JD significantly alters the pattern of inter-residue interactions within NBD of Hsp70, particularly in the catalytic site of NBD. Notably, T199, the residue critical for the ATPase activity of Hsp70,^17^ shows increased interactions with the γ-phosphate of ATP upon JD binding. This observation is in line with the previous findings, which suggested the role of threonine hydroxyl group in facilitating the ATP hydrolysis by stabilization of the transition state.^18,19,32^ Other observed changes in inter-residue interactions in lobe IIA are consistent with the recent studies on the substrate-bound Hsp70,^4^ which describe a displacement of the G loop (196-199) within a stimulating state of Hsp70 (S) preceding ATP hydrolysis, and with NMR studies describing residues within NBD most affected by the transition from ATP- to ADP-bound states of NBD.^33^ Our simulations also reproduced the movement of the Y145 residue, which accompanies the client-induced transition from the catalytically non-active restraining state (R) to the stimulating state which rapidly hydrolyses ATP (S). In the crystal structure of the S-state, the hydroxyl group of Y145 stabilizes the γ-phosphate of ATP through the interaction mediated by a water molecule located at a site created by mutating T199 to alanine. Since in our simulation T199 is restored, its hydroxyl group takes over the role of this water molecule (Figure S19). These observations pinpoint the crucial role of T199 and suggest that displacement of Y145 is a common step in the stimulation of Hsp70 by both client proteins and JD.

Based on the conformation of the T199 sidechain in respect to the γ-phosphate of ATP, we introduced two states: Thr-close, in which the hydroxyl group of T199 is in the direct contact with the γ-phosphate of ATP and the Thr-far state, in which T199 is displaced from the ATP. Our free energy profiles indicated that in Hsp70 alone Thr-far state is by 6 kcal/mol more favorable than the Thr-close state. However, the presence of JD promotes the conformational transition to the Thr-close state, making it more favorable than the Thr-far state.

To identify the allosteric mechanism by which JD promotes the Thr-close state of Hsp70, we performed additional analyses utilizing graph theory - an approach shown to provide in-depth insights into allosteric communication of biomolecules.^31,34^ Using the dynamic network analysis approach, we established that the allosteric signal from the conserved HPD motif of JD is mediated by the interdomain linker, *β*-strands 13 and 14 of NBD IIA subdomain (residues D201, I202, S203, I204, I205, E206, I207, E213, F216, L392, L391). The proposed signaling pathway passes through the residues 201-203 of Hsp70, which is supported by previously studied point-mutations (D201N, I202A, S203A), which disrupt Hsp70 stimulation by JD.^20,27^ In contrast, previously proposed pathways in the crystal structure lacking T199 only describe signaling towards K70, which plays a role in coordinating the attacking water molecule in ATP hydrolysis, and involve mainly the linker, R167, lobe IA, and IB, and SBDβ.^10^

Our results also support the contribution of JD hydrophobic and polar residues (R22, Y25, K26, F47) in stabilizing the Hsp70/JDP interface (Figure S10). Their significance in forming a stable Hsp70/JDP complex was also highlighted by the experimental work of Genevaux et al,^23^ where alanine substitutions at Y25, K26, or F47 in JD completely abolish bacterial growth. To clarify the role of specific residues within the HPD motif (residues 33–35) in the Hsp70 stimulation orchestrated by the JD, we investigated how the alanine substitution of H, P or D affects the Hsp70/JD communication. We found that the D35A substitution abolished the binding of JD to Hsp70 (Figure S12), whereas the H33A mutation did not alter the Hsp70-JD complex structure. Our free energy calculations revealed that the H33A variant’s affinity for Hsp70 is similar to that of the native JD (Figure S14A). However, additional simulations showed that despite the unaltered complex structure, this mutation prevents the transition to the Thr-close state, making it 3 kcal/mol less favorable than the Thr-far state (Figure S14; right). Unlike H33A, the D35A JD has a lower affinity for Hsp70 (2 kcal/mol less than the wild-type JD) and locates around 0.2 nm further from the NBD. In turn, substitution of the helix-breaking proline causes the reorientation of H33 and D35, disrupting their arrangement necessary for interaction with both L391 and R167 of Hsp70. Additionally, Hsp70 (R167A)-JD (D35N) variant presents similar allosteric pathways and shows similar energy profile for T199 far-close transition to wild type JD-Hsp70 complex (Figure S7, S20, S21). This is in agreement with biochemical and in vivo experiments showing partially restored Hsp70 function.^28,29^ Therefore, it can be concluded that the presence of all three amino acids is indispensable for the proper anchoring of the JD to the Hsp70 and allosteric signal transmission within Hsp70.

Finally, we investigated the conformational changes arising from the tight interaction of the HPD motif with NBD, which lead to the conformational transition of Hsp70 and T199 positioning next to the ATP γ-phosphate. We identified that adopting the Thr-close conformation is associated with the disruption of *β*-sheet conformation, formed by *β*-strands 13 and 14 within the NBD (residues 198-206; 219-227). Surprisingly, mimicking *β*-sheet disruption in the absence of JD had no effect on the Hsp70’s preference for the Thr-close state. Conversely, when JD is bound to Hsp70, restraining the *β*-sheet secondary structure prevents the transition of Hsp70 to the Thr-close state. These findings indicate that *β*-sheet disruption is a necessary, but not in itself sufficient step for the T199 - ATP γ-phosphate contact formation and further support the role of *β*-strands 13 and 14 in allosteric signal transduction.

Since inducing *β*-sheet disruption without JD failed to stimulate the Hsp70, we propose that the JD binding facilitates the transition to the previously inaccessible hydrolysis competent state (S), via an induced fit mechanism, with our work describing conformational changes associated with this process. At the molecular level, JD first forms contacts with Hsp70, promoting undocking of the NBD-SBDβ domains and forming an optimal interface between the SBDβ and NBD. The resulting push on the NBD causes the disruption of the *β*-strands 13 and 14 and subsequent movement of Thr199 towards γ-phosphate of ATP (Figure 5). Our results indicate that competition between JD helix III - SBDβ and NBD-SBDβ interfaces is necessary for partial undocking of NBD-SBDβ. Moreover, we found that JD without helix III does not favor the Thr-close state, indicating that helix III is required for β-sheet disruption and formation of the T199–ATP contact. Taken together, our results suggest a following role of JD components: (i) helix II is responsible for recognition and binding to the charged region of Hsp70 NBD; (ii) the HPD motif ensures proper orientation of JD with respect to the Hsp70, mainly through interaction with R167; (iii) helix III is necessary for partial undocking of NBD–SBDβ, promoting the conformational changes in the NBD catalytic site.

## Methods

### Simulation protocol

All simulations were performed using Gromacs 5.0.4^35^ with the Plumed 2.1 plugin.^36^ CHARMM36m force field^37^ was used for proteins, ions and Mg-ATP, and the TIP3P model^38^ was used for water. In each of the simulation boxes, the numbers of Na^+^ and Cl^−^ ions were adjusted to 0.15 M. Temperature was kept at 310 K with the v-rescale algorithm^39^ using a coupling constant of 0.1 ps. Pressure was kept at 1 bar using the Parrinello-Rahman algorithm^40^ with a coupling time of 5 ps. Periodic boundary conditions were applied and the Particle Mesh Ewald summation^41^ was used to calculate long-range electrostatic interactions with a cut-off radius of 1 nm and a Fourier grid spacing of 0.12 nm. Van der Waals interactions were calculated with Lennard-Jones potential with a cut-off radius of 1 nm. All bonds involving hydrogen were constrained using the LINCS algorithm.^42^ Leap-frog Verlet algorithm^43^ was used to integrate equations of motion with a time step of 2 fs.

The energy of the systems was minimized in two stages using steepest descent algorithm: at first, all heavy atoms of the protein were kept fixed; in the second step only the position of the protein backbone was restrained, allowing protein side chains to relax.^44,45^.After that, the systems were relaxed with short MD simulation (10 ns) with constrained protein backbone.

### Conventional MD simulations

The Hsp70 system was prepared using 4JNE PDB structure^46^ placed in a ∼13.5 nm x 13.5 nm x 13.5 nm box solvated with ∼55000 water molecules. The system was simulated in two copies and for each copy the 2.5 *µ*s-long trajectory was obtained. The Hsp70-JD system was prepared using 5NRO PDB structure^10^ placed in a ∼13 nm x 13 nm x 13 nm box solvated with ∼52000 water molecules. The system was simulated in five copies and for each copy the 1.8 *µ*s-long trajectory was obtained. In each system, the crystal water molecules coordinated to Mg^2+^ ion in Mg-ATP were kept. As the terminal region of SBD*α* is dispensable for the transmission of the allosteric signal^4,15^ to reduce the size of the system and thus, to allow for the better convergence of the free energy profiles, the C-terminal region of the lid (residues 534-609) was removed. The final structures from these simulations were used as the initial configurations for the subsequent free energy calculations.

The WT, H33A, P34A and D35A JD systems (residues 3-65) were prepared using JD from the 5NRO PDB structure.^10^ Each of the systems was placed in the ∼6 nm x 6 nm x 6 nm box, solvated with ∼7000 water molecules and simulated for 1 *µ*s.

### Trajectories analysis

Principal Component Analysis was performed for the backbone of the Hsp70’s NBD (residues 2-384) with gmx covar and gmx anaeig tools.^35^ The eigenvectors for JD-induced NBD conformational changes were assigned based on the merged NBD trajectories from Hsp70 and Hsp70-JD systems. The merged trajectory was then projected onto the first eigenvector.

Torsion angle of NBD subdomains was defined as the angle between center of mass of the backbone atoms of lobes IB (residues 40-115), IA (residues 1-39, 116-188, 361-384), IIA (residues 189-228, 307-360) and IIB (residues 229-306).^47^

Contact analysis was performed using GetContacts tool^48^ for the NBD and interdomain linker of Hsp70 (residues 2-394) using merged trajectories of Hsp70 and Hsp70-JD complex. The interactions were classified as differentiating if the difference in contact frequency between Hsp70 and Hsp70-JD systems was greater than 40 percentage points. For defining inter-residue contacts following types of interactions were considered: hydrogen bonds, ionic interactions, *π*-stacking, *π*-cation interactions and hydrophobic interactions (defined as Van der Waals interactions between purely hydrophobic residues). Default GetContacts criteria were used for all interaction types with exception of the criteria for the hydrogen bond detection were the α_DH···A_ angle value greater than or equal to 150 ° was required. Calculated contact frequencies were then visualized as chord diagrams using the Flareplot tool.^48^ NBD residues were classified as interacting with ATP if they remained in contact for at least 50% of the trajectory. Residues were considered to be at the NBD interlobular interface if they remained in contact with residues from the opposite NBD lobe for at least 50% of the trajectory.

Dynamic network analysis was performed with dynetan 1.0.1 python package.^31^ Merged trajectories of Hsp70-JD system were used to infer common network topology and perform community and betweenness centrality analyses. For tracing allosteric pathways between JD D35 and DnaK T199 weights of connections in the network were calculated using individual trajectories as separate analysis windows, while keeping previously obtained network topology. Nodes of the network were defined as the Cα atoms positions of the NBD and interdomain linker (residues 2-394) or JD (residues 3-65). Additional nodes were assigned for the magnesium ion associated with ATP molecule in the catalytic site of Hsp70, as well as for the ATP itself (two nodes, matching the positions of N1 atom of purine ring and terminal phosphorus atom). Nodes were connected by edge if any heavy atoms from two nodes remained within 0.45 nm from each other for at least 75% of the simulation length. In the case of ATP, the heavy atoms of the purine ring were assigned to the N1 atom node, whereas heavy atoms of phosphate groups and ribose were assigned to the γ-phosphate node. For each connection, the weight of the edge was calculated as the generalized correlation (GC) coefficient, which captures both linear and nonlinear correlations between residues.

Clustering of the densely interconnected nodes into communities was performed using the Louvain method,^49^ taking into the account the weight of the edges. Communities that made up for less than 1% of all the nodes were discarded in further analysis. The strength of the connection between the communities was quantified using the betweenness centrality measure, defined as the fraction of the shortest pathways between two nodes passing through a selected node, showing which nodes play a bridging role in the network.

For each of the simulated replicas, the allosteric pathways between D35 from JD conserved HPD motif (“source”) and T199 of NBD (“sink”) were calculated using Floyd-Warshall algorithm.^50^ For given edge between atoms *i* and *j*, the distance parameter *d_ij_*, defined as *d_ij_* = −log(*GC_ij_*), was calculated and the optimal path was assigned in such a way that the sum of distances along the pathway from the “source” to “sink” was minimized. Additionally, twenty sub-optimal paths were determined. Based on the calculated paths, the frequency of occurring in the path was calculated.

The antiparallel beta sheet content (*s*) of residues 198-206 and 219-227 of Hsp70 was calculated using ANTIBETARMSD method^51^ from Plumed plugin with default parameter values and strands cutoff equal to 1.

In the WT and P34A JD systems the torsion angle between H33 and D35 residues was defined as the angle between H33 N*ɛ*, H33 C*β*, D35 C*α* and D35 C*γ*.

The probability distributions for distance between Helix III of JD (residues 41-58) and SBD*β* (residues 395-505) and the distance between HPD motif and R167 were calculated using gmx distance and gmx mindist tools,^35^ respectively.

All molecular images were created using VMD and Blender.^52^ ^53^

### Free energy calculations

All of the free energy profiles were computed with the replica-exchange umbrella sampling (REUS) method^54^ and determined using the weighted histogram analysis method.^55^ Uncertainties of the free energy profiles were estimated using bootstrap error analysis taking into account the autocorrelation in the analyzed time series.

### Conformational change in the catalytic site of NBD

The free energy profiles associated with the T199 conformational change in the catalytic site of the NBD were calculated along the reaction coordinate defined as the distance between the T199 hydroxyl oxygen and the terminal phosphorus atom of ATP. The initial configurations for REUS simulations were obtained with 80 ns-long steered-MD simulation, during which the distance between T199 and ATP was gradually increased using a moving one-sided potential with a force constant of 2000 kJ · mol ^−1^ · nm ^−2^. Ten equally spaced US windows were used, spanning the range of 0.3 to 1.2 nm of the reaction coordinate. The spring constant of the harmonic biasing potential was set to 1500 kJ · mol ^−1^ · nm ^−2^ and each of the REUS windows was simulated for 1 *µ*s. The same procedure was applied for the substitution variants.

Additionally, during the investigation of the beta-sheet disruption influence on the Thr-far to Thr-close transition additional simulations were performed. In the Hsp70-JD system, the beta-sheet was restrained by setting the parameter *s* to 2.73, whereas in the Hsp70 system the beta-sheet was disrupted by keeping the *s* parameter at the value of 1.8. In both cases the one-sided harmonic potential was used, with the spring constant of 4000 kJ · mol ^−1^ · nm ^−2^.

In the case of the ΔH3-4 system, during the free energy calculations the conformation of the remaining JD (residues 3-40) was restrained with the additional harmonic biasing potential (with the spring constant set to 5000 kJ · mol ^−1^ · nm ^−2^), which kept the 0.0 nm RMSD value with respect to the wild-type helix II conformation.

### JD positioning on Hsp70

The free energy profiles associated with the JD binding on Hsp70 were calculated along the reaction coordinate defined as the distance between the center of mass of backbone atoms of the JD (residues 25-31 and 48-54) and the center of mass of C*α* atoms of Hsp70 binding site (residues 208-216 and 415-422). The initial configurations for REUS simulations were obtained with 100 ns-long steered-MD simulation, during which the distance between Hsp70 and JD was gradually increased using a moving one-sided harmonic potential with a force constant of 1500 kJ · mol ^−1^ · nm ^−2^. Six equally spaced US windows were used, spanning the range of 0.8 to 1.3 nm of the reaction coordinate. The spring constant of the harmonic biasing potential was set to 3500 kJ · mol ^−1^ · nm ^−2^ and each of the REUS windows was simulated for 800 ns.

### SBD and NBD dissociation

In the studies of SBD and NBD dissociation the reaction coordinate was defined as the distance between the center of mass of C*α* atoms of the SBD*β* domain (residues 395-505) and the center of mass of C*α* atoms of the NBD fragments, which are in direct contact with SBD*β* (residues 146-226). To make the system significantly smaller and thus, to achieve better convergence of the free energy profiles, the lid sub-domain, SBD*α*, was truncated by removing residues 506-602 of Hsp70.

The initial configurations for REUS simulations were obtained with 500 ns-long steered MD simulations, during which the distance between NBD and SBD*β* was gradually increased using a moving one-sided potential with a spring constant of 2500 kJ · mol ^−1^ · nm ^−2^, from the initial value of 2.58 nm up to 3.58 nm. Five equally spaced US windows were used, spanning the range of 2.5 to 2.9 nm of the reaction coordinate. The spring constant of the harmonic biasing potential was set to 3500 kJ · mol ^−1^ · nm ^−2^ and each of the REUS windows was simulated for 1 *µ*s.

## Acknowledgement

This study was supported by National Science Center, Poland (OPUS 21 2021/41/B/NZ8/02835) (to BT). We gratefully acknowledge Poland’s high-performance Infrastructure PLGrid ACK Cyfronet AGH, PCSS, for providing computer facilities and support within computational grant no PLG/2022/015642 and PLG/2025/017999. We thank Dr. Agnieszka Kłosowska for helpful discussions.

## References

(1) Richter, K.; Haslbeck, M.; Buchner, J. The Heat Shock Response: Life on the Verge of Death. Mol. Cell 2010, 40 (2), 253–266. 10.1016/j.molcel.2010.10.006.

(2) Kityk, R.; Kopp, J.; Sinning, I.; Mayer, M. P. Structure and Dynamics of the ATP-Bound Open Conformation of Hsp70 Chaperones. Mol. Cell 2012, 48 (6), 863–874. 10.1016/j.molcel.2012.09.023.

(3) Laufen, T.; Mayer, M. P.; Beisel, C.; Klostermeier, D.; Mogk, A.; Reinstein, J.; Bukau, B. Mechanism of Regulation of Hsp70 Chaperones by DnaJ Cochaperones. Proc. Natl. Acad. Sci. 1999, 96 (10), 5452–5457. 10.1073/pnas.96.10.5452.

(4) Wang, W.; Liu, Q.; Liu, Q.; Hendrickson, W. A. Conformational Equilibria in Allosteric Control of Hsp70 Chaperones. Mol. Cell 2021, 81 (19), 3919–3933.e7. 10.1016/j.molcel.2021.07.039.

(5) Kityk, R.; Vogel, M.; Schlecht, R.; Bukau, B.; Mayer, M. P. Pathways of Allosteric Regulation in Hsp70 Chaperones. Nat. Commun. 2015, 6 (8308). 10.1038/ncomms9308.

(6) Misselwitz, B.; Staeck, O.; Rapoport, T. A. J Proteins Catalytically Activate Hsp70 Molecules to Trap a Wide Range of Peptide Sequences. Mol. Cell 1998, 2 (5), 593–603. 10.1016/S1097-2765(00)80158-6.

(7) Mayer, M. P.; Schröder, H.; Rüdiger, S.; Paal, K.; Laufen, T.; Bukau, B. Multistep Mechanism of Substrate Binding Determines Chaperone Activity of Hsp70. Nat. Struct. Biol. 2000, 7 (7), 586–593. 10.1038/76819.

(8) Harrison, C. J.; Hayer-Hartl, M.; Liberto, M. D.; Hartl, F.-U.; Kuriyan, J. Crystal Structure of the Nucleotide Exchange Factor GrpE Bound to the ATPase Domain of the Molecular Chaperone DnaK. Science 1997, 276 (5311), 431–435. 10.1126/science.276.5311.431.

(9) Liu, Q.; Liang, C.; Zhou, L. Structural and Functional Analysis of the Hsp70/Hsp40 Chaperone System. Protein Sci. 2020, 29 (2), 378–390. 10.1002/pro.3725.

(10) Kityk, R.; Kopp, J.; Mayer, M. P. Molecular Mechanism of J-Domain-Triggered ATP Hydrolysis by Hsp70 Chaperones. Mol. Cell 2018, 69 (2), 227–237.e4. 10.1016/j.molcel.2017.12.003.

(11) Astl, L.; Verkhivker, G. M. Dynamic View of Allosteric Regulation in the Hsp70 Chaperones by J-Domain Cochaperone and Post-Translational Modifications: Computational Analysis of Hsp70 Mechanisms by Exploring Conformational Landscapes and Residue Interaction Networks. J. Chem. Inf. Model. 2020, 60 (3), 1614–1631. 10.1021/acs.jcim.9b01045.

(12) Tomiczek, B.; Delewski, W.; Nierzwicki, L.; Stolarska, M.; Grochowina, I.; Schilke, B.; Dutkiewicz, R.; Uzarska, M. A.; Ciesielski, S. J.; Czub, J.; Craig, E. A.; Marszalek, J. Two-Step Mechanism of J-Domain Action in Driving Hsp70 Function. PLOS Comput. Biol. 2020, 16 (6), 1–29. 10.1371/journal.pcbi.1007913.

(13) Karzai, A. W.; McMacken, R. A Bipartite Signaling Mechanism Involved in DnaJ-Mediated Activation of the Escherichia Coli DnaK Protein. J. Biol. Chem. 1996, 271 (19), 11236–11246. 10.1074/jbc.271.19.11236.

(14) Delewski, W.; Paterkiewicz, B.; Manicki, M.; Schilke, B.; Tomiczek, B.; Ciesielski, S. J.; Nierzwicki, L.; Czub, J.; Dutkiewicz, R.; Craig, E. A.; Marszalek, J. Iron–Sulfur Cluster Biogenesis Chaperones: Evidence for Emergence of Mutational Robustness of a Highly Specific Protein–Protein Interaction. Mol. Biol. Evol. 2015, 33 (3), 643–656.

(15) Bascos, N. A. D.; Mayer, M. P.; Bukau, B.; Landry, S. J. The Hsp40 J-Domain Modulates Hsp70 Conformation and ATPase Activity with a Semi-Elliptical Spring. Protein Sci. 2017, 26 (9), 1838–1851. 10.1002/pro.3223.

(16) Hendrickson, W. A. Theory of Allosteric Regulation in Hsp70 Molecular Chaperones. QRB Discov. 2020, 1, e7. 10.1017/qrd.2020.10.

(17) Barthel, T. K.; Zhang, J.; Walker, G. C. ATPase-Defective Derivatives of Escherichia Coli DnaK That Behave Differently with Respect to ATP-Induced Conformational Change and Peptide Release. J. Bacteriol. 2001, 183 (19), 5482–5490. 10.1128/JB.183.19.5482-5490.2001.

(18) Kiani, F. A.; Fischer, S. Comparing the Catalytic Strategy of ATP Hydrolysis in Biomolecular Motors. Phys Chem Chem Phys 2016, 18 (30), 20219–20233. 10.1039/C6CP01364C.

(19) Suh, W.-C.; Burkholder, W. F.; Lu, C. Z.; Zhao, X.; Gottesman, M. E.; Gross, C. A.; Prieß, M.; Göddeke, H.; Groenhof, G.; Schäfer, L. V. Molecular Mechanism of ATP Hydrolysis in an ABC Transporter. Proc. Natl. Acad. Sci. 2018, 4 (10), 1334–1343. 10.1021/acscentsci.8b00369.

(20) Chang, L.; Thompson, A. D.; Ung, P.; Carlson, H. A.; Gestwicki, J. E. Mutagenesis Reveals the Complex Relationships between ATPase Rate and the Chaperone Activities of Escherichia Coli Heat Shock Protein 70 (Hsp70/DnaK). J. Biol. Chem. 2010, 285 (28), 21282–21291. 10.1074/jbc.M110.124149.

(21) Schneider, M.; Antes, I. Comparison of Allosteric Signaling in DnaK and BiP Using Mutual Information between Simulated Residue Conformations. Proteins Struct. Funct. Bioinforma. 2023, 91 (2), 237–255. 10.1002/prot.26425.

(22) Wall, D.; Zylicz, M.; Georgopoulos, C. The NH2-Terminal 108 Amino Acids of the Escherichia Coli DnaJ Protein Stimulate the ATPase Activity of DnaK and Are Sufficient for Lambda Replication. J. Biol. Chem. 1994, 269 (7), 5446–5451. 10.1016/S0021-9258(17)37706-2.

(23) Genevaux, P.; Schwager, F.; Georgopoulos, C.; Kelley, W. L. Scanning Mutagenesis Identifies Amino Acid Residues Essential for the in Vivo Activity of the Escherichia Coli DnaJ (Hsp40) J-Domain. Genetics 2002, 162 (3), 1045–1053. 10.1093/genetics/162.3.1045.

(24) Li, S. C.; Goto, N. K.; Williams, K. A.; Deber, C. M. Alpha-Helical, but Not Beta-Sheet, Propensity of Proline Is Determined by Peptide Environment. Proc. Natl. Acad. Sci. 1996, 93 (13), 6676–6681. 10.1073/pnas.93.13.6676.

(25) English, C. A.; Sherman, W.; Meng, W.; Gierasch, L. M. The Hsp70 Interdomain Linker Is a Dynamic Switch That Enables Allosteric Communication between Two Structured Domains. J. Biol. Chem. 2017, 292 (36), 14765–14774. 10.1074/jbc.M117.789313.

(26) Szyperski, T.; Pellecchia, M.; Wall, D.; Georgopoulos, C.; Wüthrich, K. NMR Structure Determination of the Escherichia Coli DnaJ Molecular Chaperone: Secondary Structure and Backbone Fold of the N-Terminal Region (Residues 2-108) Containing the Highly Conserved J Domain. Proc. Natl. Acad. Sci. 1994, 91 (24), 11343–11347. 10.1073/pnas.91.24.11343.

(27) Kamath-Loeb, A. S.; Lu, C. Z.; Suh, W.-C.; Lonetto, M. A.; Gross, C. A. Analysis of Three DnaK Mutant Proteins Suggests That Progression through the ATPase Cycle Requires Conformational Changes (∗). J. Biol. Chem. 1995, 270 (50), 30051–30059. 10.1074/jbc.270.50.30051.

(28) Landry, S. J. Structure and Energetics of an Allele-Specific Genetic Interaction between dnaJ and dnaK: Correlation of Nuclear Magnetic Resonance Chemical Shift Perturbations in the J-Domain of Hsp40/DnaJ with Binding Affinity for the ATPase Domain of Hsp70/DnaK. Biochemistry 2003, 42 (17), 4926–4936. 10.1021/bi027070y.

(29) Suh, W.-C.; Burkholder, W. F.; Lu, C. Z.; Zhao, X.; Gottesman, M. E.; Gross, C. A. Interaction of the Hsp70 Molecular Chaperone, DnaK, with Its Cochaperone DnaJ. Proc. Natl. Acad. Sci. 1998, 95 (26), 15223–15228. 10.1073/pnas.95.26.15223.

(30) Sethi, A.; Eargle, J.; Black, A. A.; Luthey-Schulten, Z. Dynamical Networks in tRNA:Protein Complexes. Proc. Natl. Acad. Sci. 2009, 106 (16), 6620–6625. 10.1073/pnas.0810961106.

(31) Melo, M. C. R.; Bernardi, R. C.; de la Fuente-Nunez, C.; Luthey-Schulten, Z. Generalized Correlation-Based Dynamical Network Analysis: A New High-Performance Approach for Identifying Allosteric Communications in Molecular Dynamics Trajectories. J. Chem. Phys. 2020, 153 (13), 134104. 10.1063/5.0018980.

(32) O’Brien, M. C.; Flaherty, K. M.; McKay, D. B. Lysine 71 of the Chaperone Protein Hsc70 Is Essential for ATP Hydrolysis. J. Biol. Chem. 1996, 271 (27), 15874–15878. 10.1074/jbc.271.27.15874.

(33) Meng, W.; Clerico, E. M.; McArthur, N.; Gierasch, L. M. Allosteric Landscapes of Eukaryotic Cytoplasmic Hsp70s Are Shaped by Evolutionary Tuning of Key Interfaces. Proc. Natl. Acad. Sci. 2018, 115 (47), 11970–11975. 10.1073/pnas.1811105115.

(34) Patel, A. C.; Sinha, S.; Palermo, G. Graph Theory Approaches for Molecular Dynamics Simulations. Q. Rev. Biophys. 2024, 57, e15. 10.1017/S0033583524000143.

(35) Van Der Spoel, D.; Lindahl, E.; Hess, B.; Groenhof, G.; Mark, A. E.; Berendsen, H. J. C. GROMACS: Fast, Flexible, and Free. J. Comput. Chem. 2005, 26 (16), 1701–1718. 10.1002/jcc.20291.

(36) Tribello, G. A.; Bonomi, M.; Branduardi, D.; Camilloni, C.; Bussi, G. PLUMED 2: New Feathers for an Old Bird. Comput. Phys. Commun. 2014, 185 (2), 604–613. 10.1016/j.cpc.2013.09.018.

(37) Huang, J.; Rauscher, S.; Nawrocki, G.; Ran, T.; Feig, M.; De Groot, B. L.; Grubmüller, H.; MacKerell, A. D. CHARMM36m: An Improved Force Field for Folded and Intrinsically Disordered Proteins. Nat. Methods 2016, 14 (1), 71–73. 10.1038/nmeth.4067.

(38) Jorgensen, W. L.; Chandrasekhar, J.; Madura, J. D.; Impey, R. W.; Klein, M. L. Comparison of Simple Potential Functions for Simulating Liquid Water. J. Chem. Phys. 1983, 79 (2), 926– 935. 10.1063/1.445869.

(39) Bussi, G.; Donadio, D.; Parrinello, M. Canonical Sampling through Velocity Rescaling. J. Chem. Phys. 2007, 126 (1). 10.1063/1.2408420.

(40) Parrinello, M.; Rahman, A. Polymorphic Transitions in Single Crystals: A New Molecular Dynamics Method. J. Appl. Phys. 1981, 52 (12), 7182–7190. 10.1063/1.328693.

(41) Essmann, U.; Perera, L.; Berkowitz, M. L.; Darden, T.; Lee, H.; Pedersen, L. G. A Smooth Particle Mesh Ewald Method. J. Chem. Phys. 1995, 103 (19), 8577–8593. 10.1063/1.470117.

(42) Hess, B.; Bekker, H.; Berendsen, H. J. C.; Fraaije, J. G. E. M. LINCS: A Linear Constraint Solver for Molecular Simulations. J. Comput. Chem. 1997, 18 (12), 1463–1472. 10.1002/(SICI)1096-987X(199709)18:12<1463::AID-JCC4>3.0.CO;2-H.

(43) van Gunsteren, W. F.; Berendsend, H. J. C. A Leap Frog Algorithm for Stochastic Dynamics. Mol. Simul. 1988, 1 (May 2013), 173–185. 10.1080/08927028808080941.

(44) Lee, J.; Cheng, X.; Swails, J. M.; Yeom, M. S.; Eastman, P. K.; Lemkul, J. A.; Wei, S.; Buckner, J.; Jeong, J. C.; Qi, Y.; Jo, S.; Pande, V. S.; Case, D. A.; Brooks, C. L. I.; MacKerell, A. D. Jr.; Klauda, J. B.; Im, W. CHARMM-GUI Input Generator for NAMD, GROMACS, AMBER, OpenMM, and CHARMM/OpenMM Simulations Using the CHARMM36 Additive Force Field. J. Chem. Theory Comput. 2016, 12 (1), 405–413. 10.1021/acs.jctc.5b00935.

(45) Jo, S.; Kim, T.; Iyer, V. G.; Im, W. CHARMM-GUI: A Web-Based Graphical User Interface for CHARMM. J. Comput. Chem. 2008, 29 (11), 1859–1865. 10.1002/jcc.20945.

(46) Qi, R.; Sarbeng, E. B.; Liu, Q.; Le, K. Q.; Xu, X.; Xu, H.; Yang, J.; Wong, J. L.; Vorvis, C.; Hendrickson, W. A.; Zhou, L.; Liu, Q. Allosteric Opening of the Polypeptide-Binding Site When an Hsp70 Binds ATP. Nat. Struct. Mol. Biol. 2013, 20 (7), 900–907. 10.1038/nsmb.2583.

(47) Arakawa, A.; Handa, N.; Shirouzu, M.; Yokoyama, S. Biochemical and Structural Studies on the High Affinity of Hsp70 for ADP. Protein Sci. 2011, 20.

(48) Venkatakrishnan, A. J.; Fonseca, R.; Ma, A. K.; Hollingsworth, S. A.; Chemparathy, A.; Hilger, D.; Kooistra, A. J.; Ahmari, R.; Babu, M. M.; Kobilka, B. K.; Dror, R. O. Uncovering Patterns of Atomic Interactions in Static and Dynamic Structures of Proteins. Biophysics November 13, 2019. 10.1101/840694.

(49) Blondel, V. D.; Guillaume, J.-L.; Lambiotte, R.; Lefebvre, E. Fast Unfolding of Communities in Large Networks. J. Stat. Mech. Theory Exp. 2008, 2008 (10), P10008. 10.1088/1742-5468/2008/10/P10008.

(50) Floyd, R. W. Algorithm 97: Shortest Path. Commun ACM 1962, 5 (6), 345. 10.1145/367766.368168.

(51) Pietrucci, F.; Laio, A. A Collective Variable for the Efficient Exploration of Protein Beta-Sheet Structures: Application to SH3 and GB1. J. Chem. Theory Comput. 2009, 5 (9), 2197– 2201. 10.1021/ct900202f.

(52) Humphrey, W.; Dalke, A.; Schulten, K. VMD: Visual Molecular Dynamics. J. Mol. Graph. 1996, 14 (1), 33–38. 10.1016/0263-7855(96)00018-5.

(53) Community, B. O. Blender - a 3D Modelling and Rendering Package; Blender Foundation: Stichting Blender Foundation, Amsterdam, 2018.

(54) Torrie, G. M.; Valleau, J. P. Nonphysical Sampling Distributions in Monte Carlo Free-Energy Estimation: Umbrella Sampling. J. Comput. Phys. 1977, 23 (2), 187–199. 10.1016/0021-9991(77)90121-8.

(55) Kumar, S.; Rosenberg, J. M.; Bouzida, D.; Swendsen, R. H.; Kollman, P. A. THE Weighted Histogram Analysis Method for Free-Energy Calculations on Biomolecules. I. The Method. J. Comput. Chem. 1992, 13 (8), 1011–1021. 10.1002/jcc.540130812.

