## Supplementary information for "Mechanism and energetics of JDP induced Hsp70’s conformational transition towards catalytically active state"

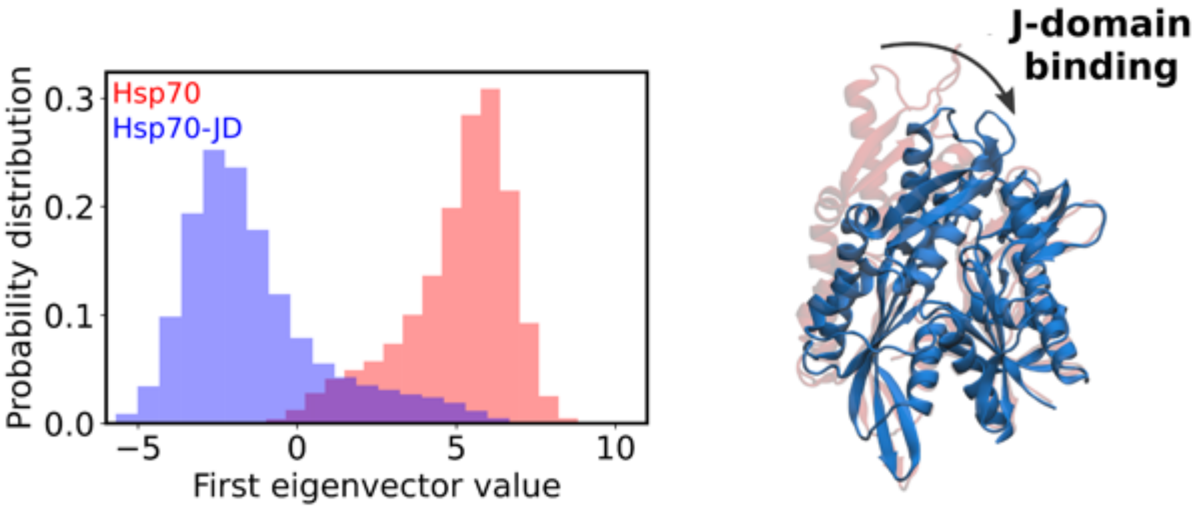


Figure S1 Distribution of the Hsp70 and Hsp70-JD trajectories projected onto the first eigenvector obtained by the Principal Component Analysis. The visual representation of the NBD along the first eigenvector is depicted on the right.

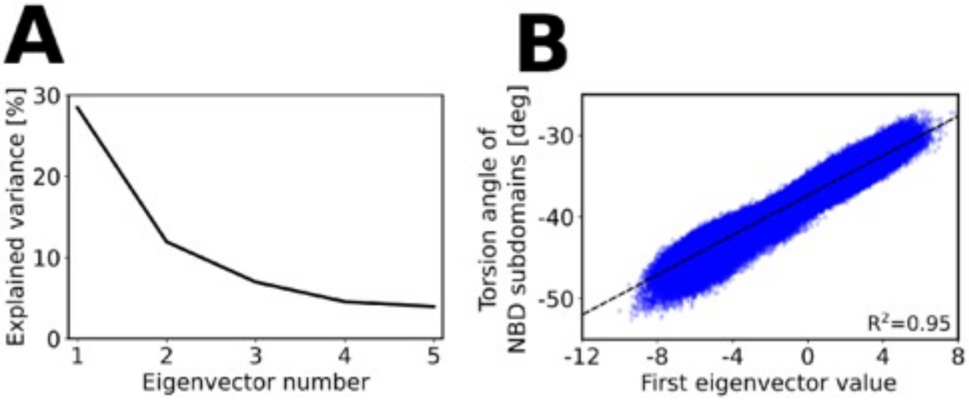


Figure S2: Results of the Principal Component Analysis for the merged Hsp70 and Hsp70-JD trajectories. (A) Percentage of the total variance explained by each of the first five eigenvectors. (B) Correlation between the values of the first component and the torsion angle of the NBD subdomains.

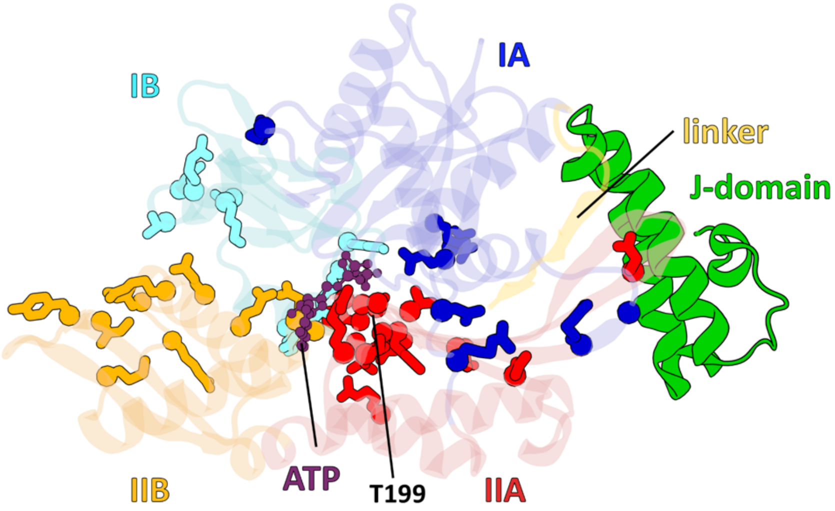


Figure S3: Residues of Hsp70 which display the most prominent differences in the contact frequency within NBD and inter-domain linker in the presence or absence of the JD. Color scheme: purple– ATP molecule, blue– lobe IA of NBD, cyan– lobe IB of NBD, red– lobe IIA of NBD, orange– lobe IIB of NBD, yellow– interdomain linker. Only NBD of DnaK showed for clarity.

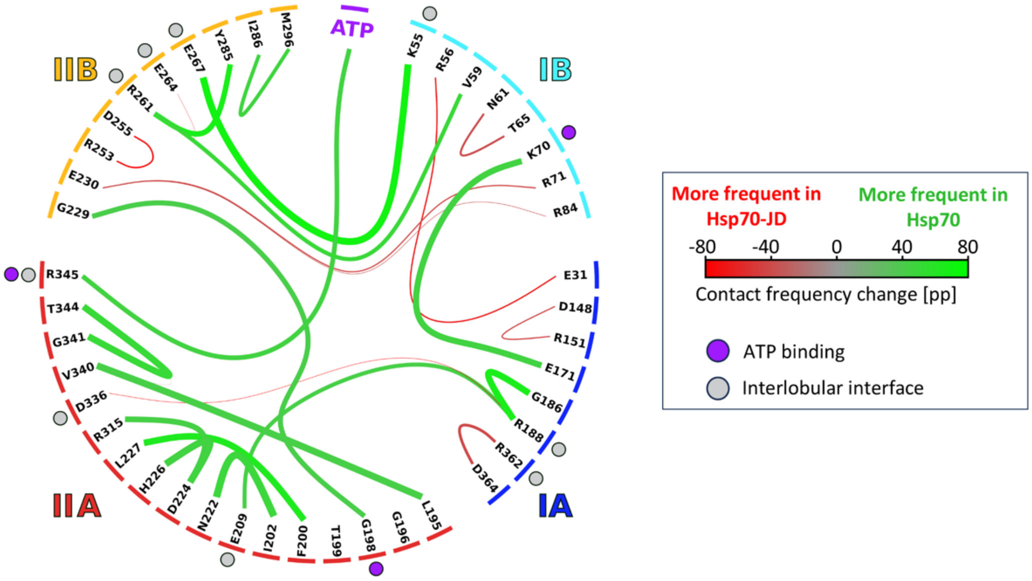


Figure S4: Chord diagram showing changes in NBD residues contact frequency associated with Hsp70 alone. Line widths are proportional to the contact frequency in Hsp70 alone and are colored based on the contact frequency difference with respect to Hsp70-JD complex. Only contacts displaying frequency difference of 40 percentage points or greater are shown. Dots positioned next to NBD residues indicate ATP-binding residues (violet) or residues positioned at the interlobular interface (gray) in the Hsp70 alone. Colored bars along the circle's rim indicate NBD subdomain containing a given residue.

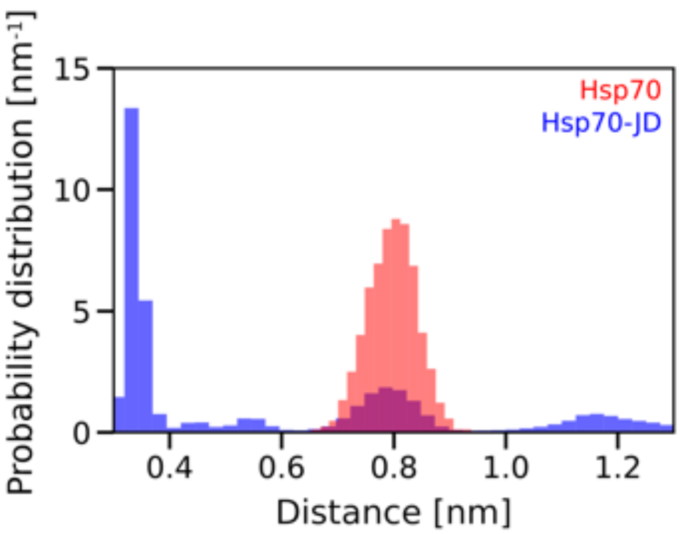


Figure S5: Probability distribution of the distance between oxygen from the hydroxyl group of T199 and the γ-phosphate of ATP during conventional MD simulations of the Hsp70 (red) and Hsp70-JD complex (blue).

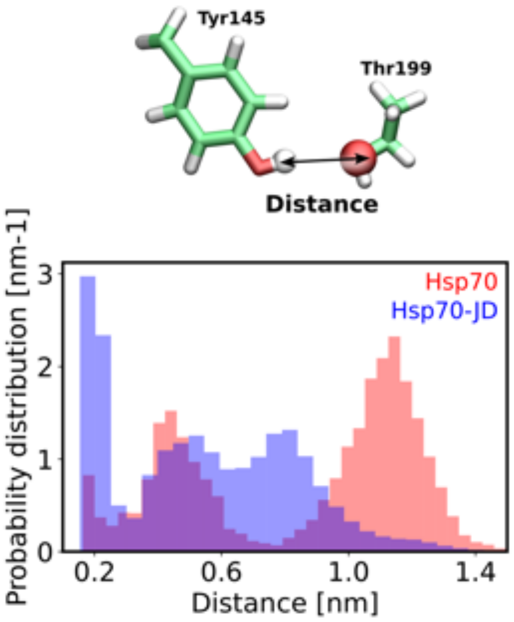


Figure S6: Probability distribution of the distance between hydrogen from the hydroxyl group of Y145 and the oxygen from the hydroxyl group of T199 of Hsp70 during conventional MD simulations of the Hsp70 (red) and Hsp70-JD complex (blue).

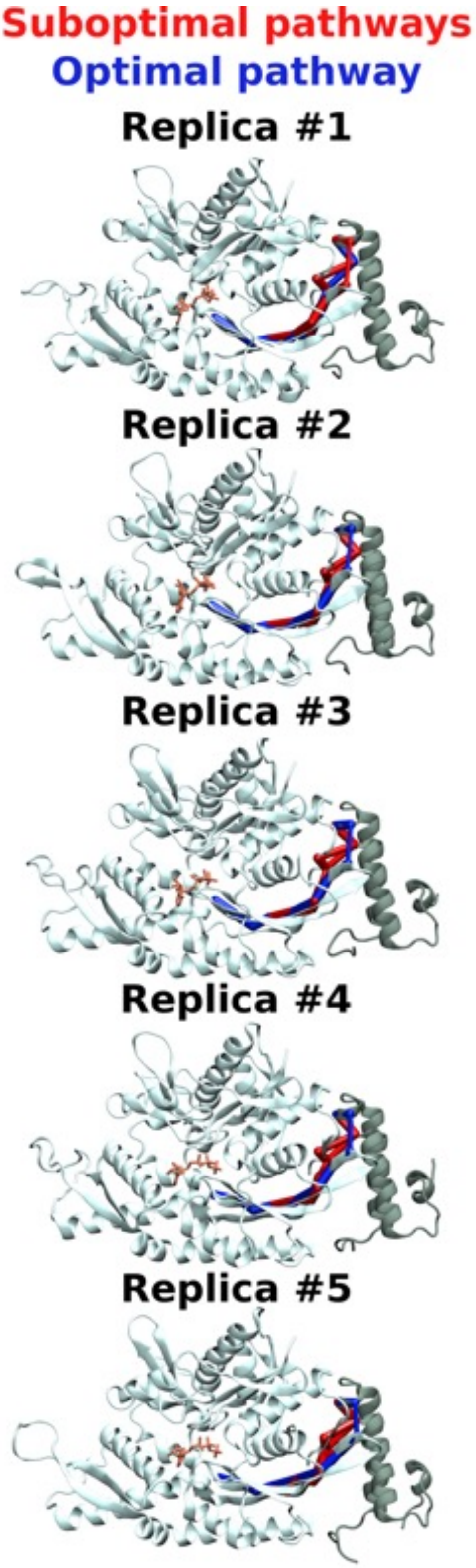


Figure S7: Allosteric pathways determined for each of the simulated replica of Hsp70-JD system. The optimal pathway was indicated in blue, whereas suboptimal pathways were depicted in red. Only NBD showed for clarity.

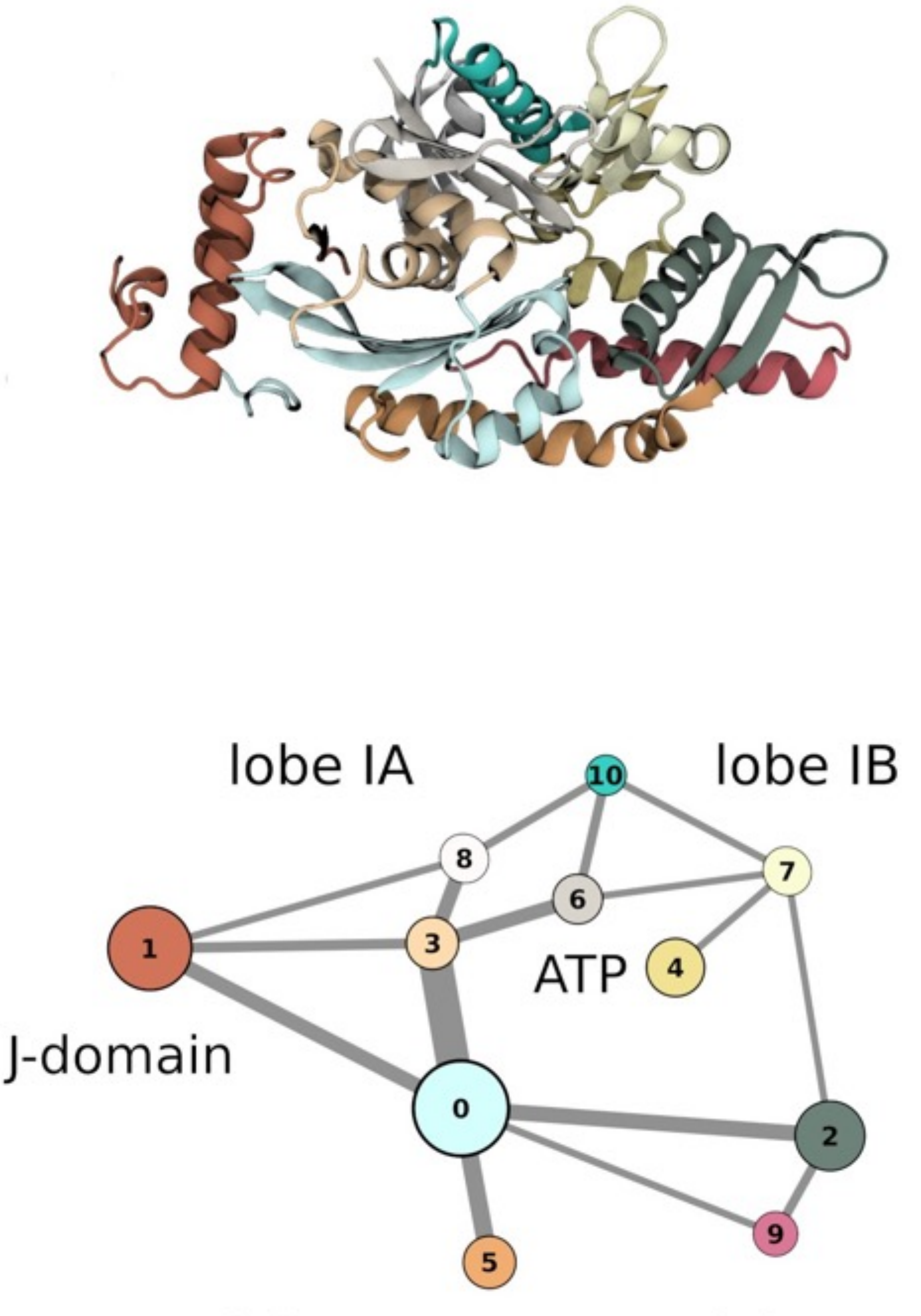


Figure S8: Structural (top) and graph (bottom) representation of the community analysis for the Hsp70-JD complex. The width of the connections between communities in the bottom panel corresponds to the betweenness centrality measure between them (for exact values see Table S1). Communities are numbered from 0 to 10. The ATP atoms were part of the community number 4 and 6.

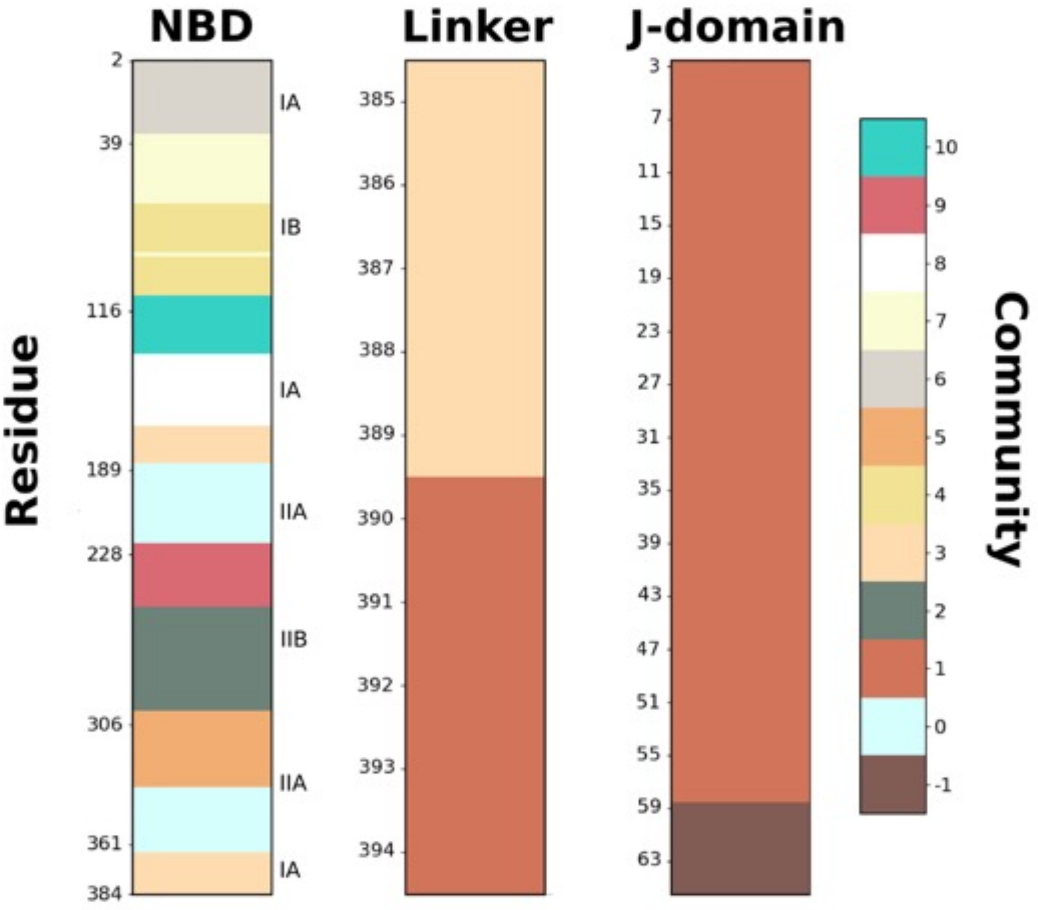


Figure S9: Composition of the communities obtained during the community network analysis of the Hsp70-JD complex. In the case of NBD, the lobes of NBD, which correspond to each residue range were indicated on the right. The -1 community represents nodes which were not assigned to any community.

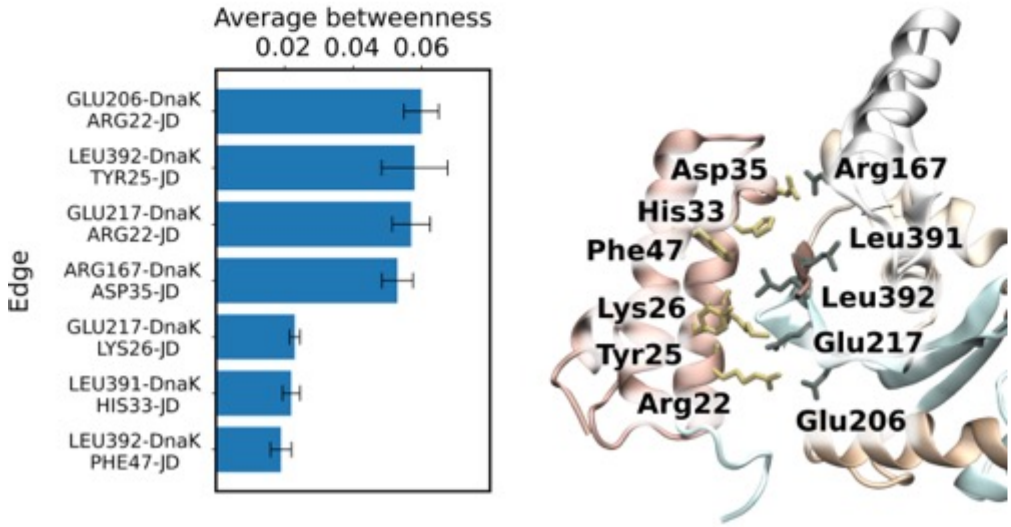


Figure S10: Edges matching the highest betweenness centrality on the interface formed between nodes of NBD and JD. The joint network was used to choose ten edges with the highest betweenness value from each of the simulated copies of the Hsp70-JD system. Only edges with betweenness value greater than 0.01 are shown. Color scheme of the proteins interface on the right corresponds to the communities composition in Figure S10.

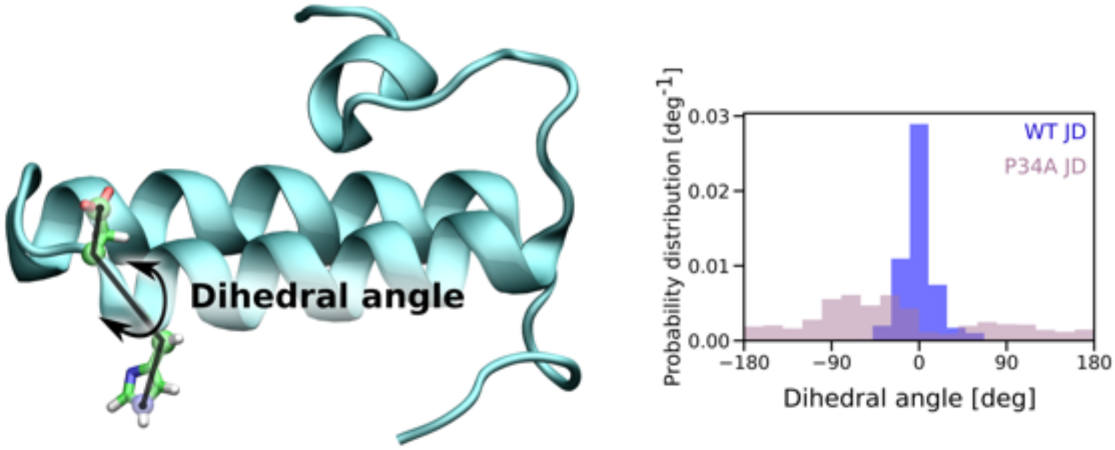


Figure S11: Probability distribution of the dihedral angle between H33 and D35 in the wild-type JD (blue) and the P34A variant (violet) of JD.

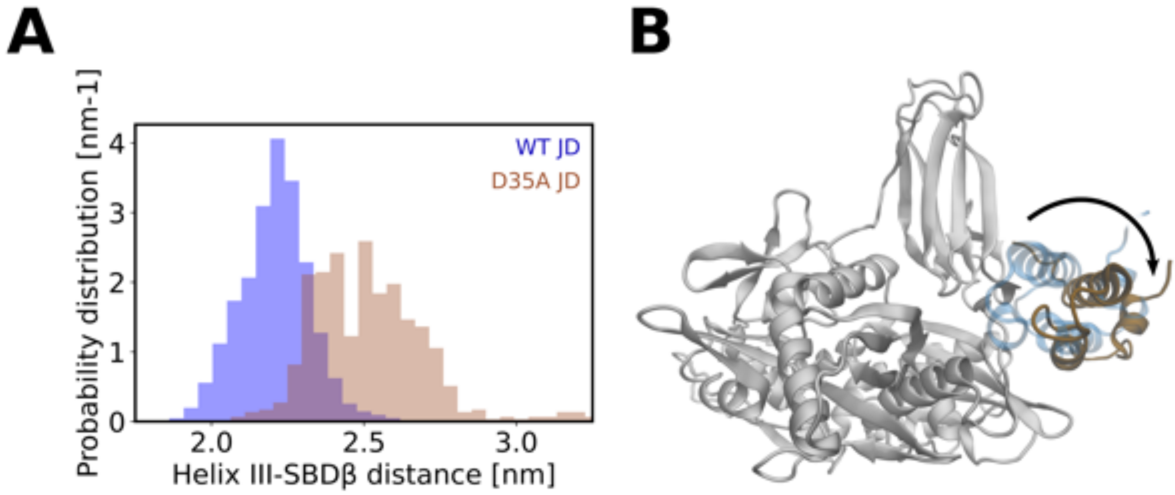


Figure S12: Dissociation of the D35A mutant of JD from the specific binding site. (A) Probability distribution of the distance between SBD$\beta$ and helix III of D35A variant (brown) and the wild-type (blue) JD (B) Position of the D35A variant of the JD bound to Hsp70 (brown) with respect to the wild-type bound JD (transparent blue).

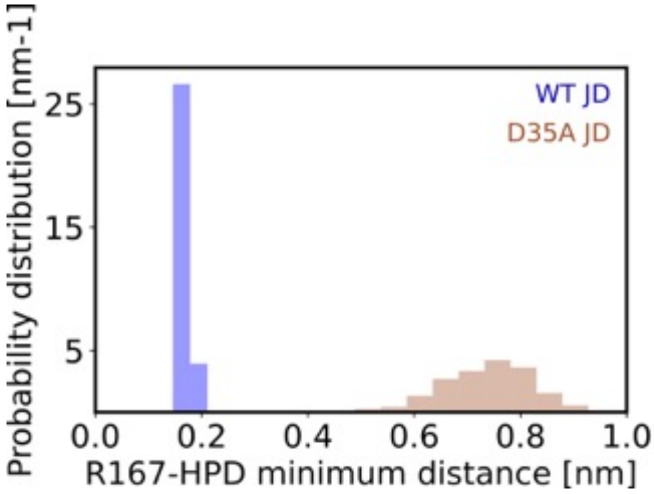

Figure S13: Probability distribution of the minimum distance between R167 of Hsp70 and the HPD motif of JD for wild-type (blue) and D35A mutant (brown).

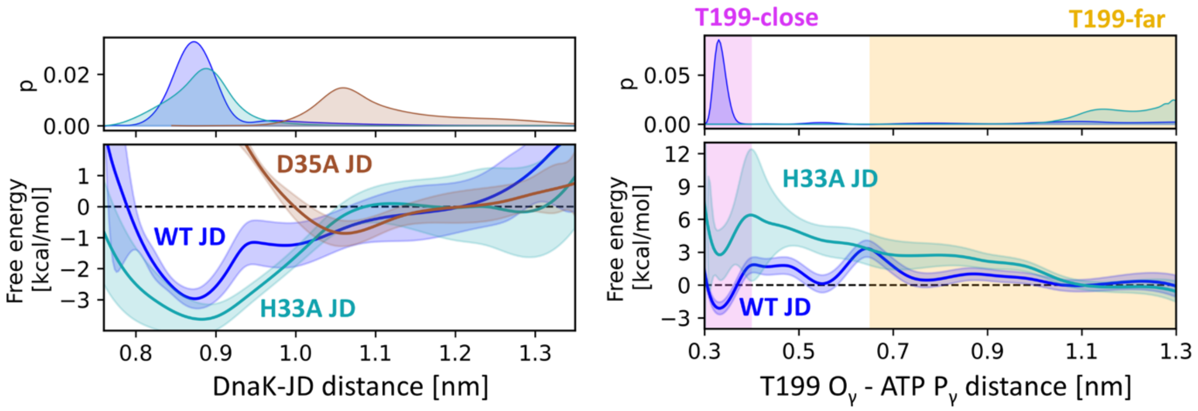


Figure S14: (Left) The probability distributions (top) and free energy profiles (bottom) for the JD association to the Hsp70. The profile for the H33A mutant was shown in cyan, profile for the D35A mutant was shown in brown, whereas profile for the wild-type JD binding was shown in blue. (Right) The probability distributions (top) and free energy profiles (bottom) for the Thr-far to Thr-close conformational change for the complex of H33A mutant (cyan) or the wild-type (blue) JD bound to Hsp70.

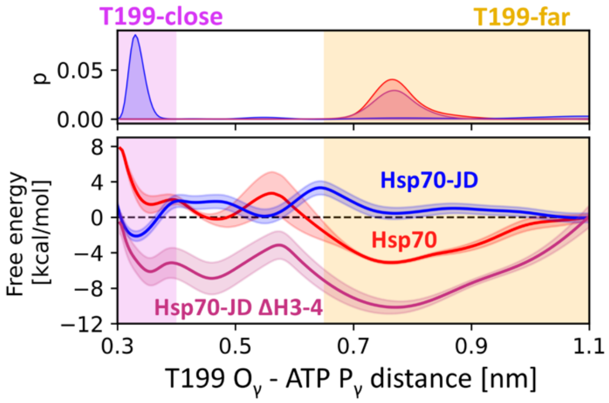


Figure S15: The probability distributions (top) and free energy profiles (bottom) for the Thr-far to Thr-close conformational change for the ΔH3-4 JD (magenta). Profiles for the Hsp70 (blue) and Hsp70-JD (red) were added for comparison.

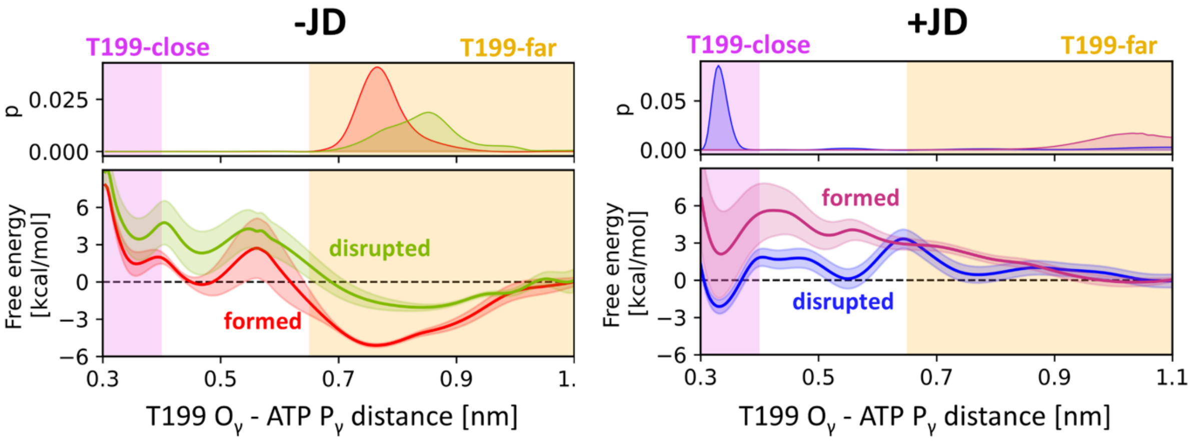


Figure S16: (Left) The probability distributions (top) and free energy profiles (bottom) for the Thr-far to Thr-close conformational change in the absence of the JD. The non-biased profile for the formed $\beta$-sheet is shown in red, whereas the profile for the bias-induced disrupted $\beta$-sheet is shown in green. (Right) The probability distributions (top) and free energy profiles (bottom) for the Thr-far to Thr-close conformational change in the presence of the JD. The non-biased profile for the disrupted $\beta$-sheet is shown in blue, whereas the profile for the bias-induced $\beta$-sheet kept is shown in magenta.

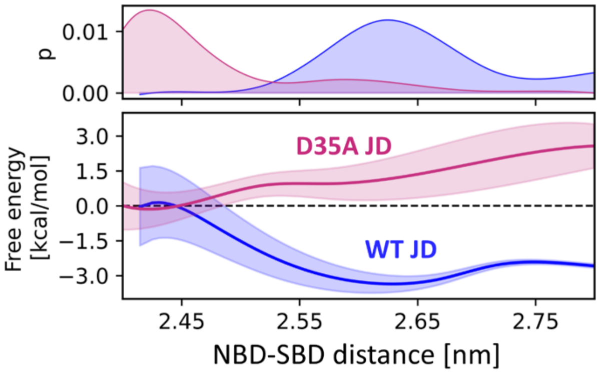


Figure S17: Probability distributions (top) and free energy profiles (bottom) for the dissociation of the NBD and SBD$\beta$ subdomains of Hsp70 caused by wild-type JD (blue) and D35A variant of JD (magenta).

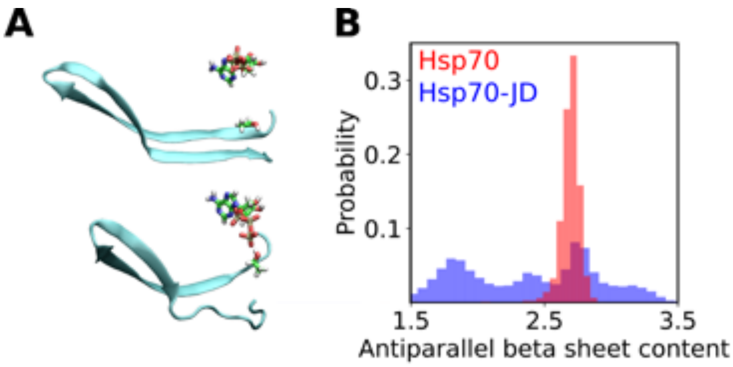

Figure S18: Disruption of the beta-sheet caused by the JD binding. (A) Structural representation of the movement of T199 towards the ATP enabled by beta-sheet disruption. The ATP and T199 are shown in licorice representation, whereas beta-sheet is shown in cartoon representation. (B) Probability distribution of the antiparallel beta-sheet content of the 198-206 and 219-227 residues of Hsp70 in the Hsp70 alone (red) and the Hsp70-JD complex (blue).

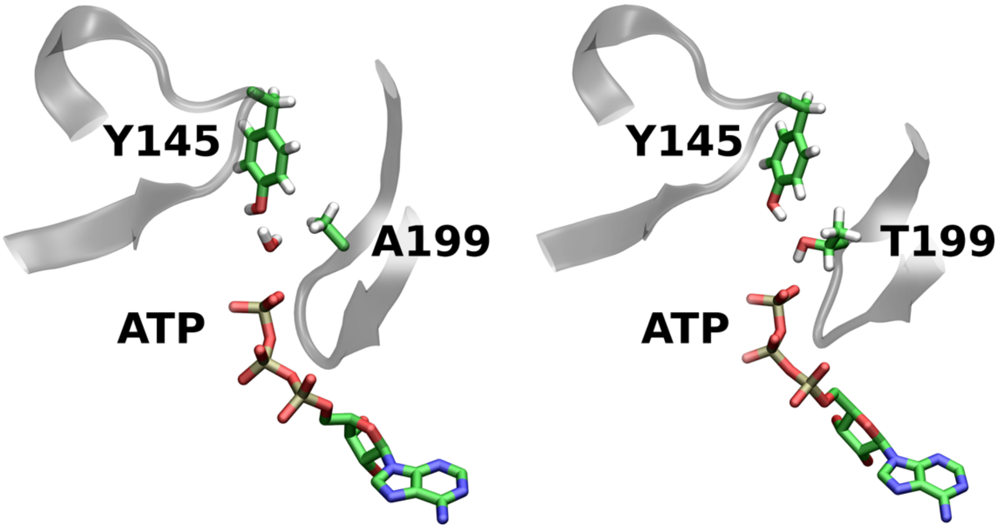


Figure S19: Coordination of the γ-phosphate of ATP in the crystal structure of the substrate-induced stimulated-state of Hsp70 (PDB id: 7KRU, left) and the Thr-close state of Hsp70 visible in our simulations of the Hsp70 with JD (right).

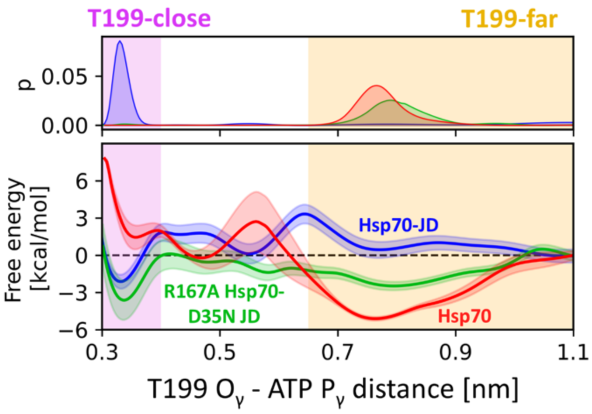


Figure S20: Probability distribution (top) and free energy profile (bottom) of the Thr-far to Thr-close conformational change for the Hsp70 (R167A) - JD (D35N) double mutant (green). Profiles for wildtype complex (blue) and Hsp70 alone (red) added for comparison.

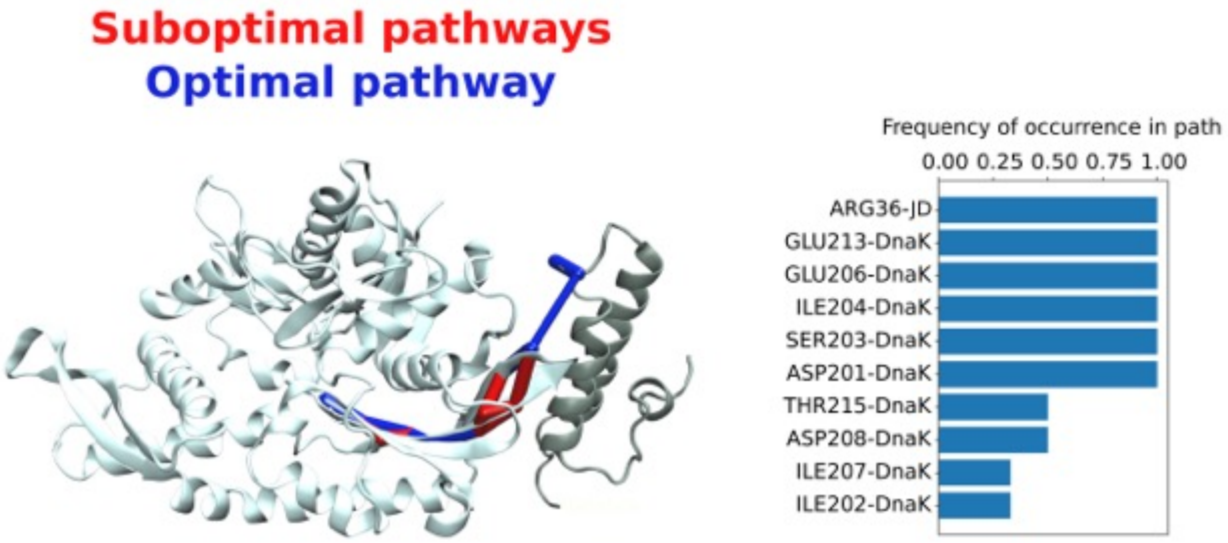


Figure S21: Allosteric pathways determined for the Hsp70 (R167A) - JD (D35N) double mutant. The pathways between HPD motif of JD and the catalytic pocket of Hsp70 were calculated from the Umbrella Sampling trajectories corresponding to the Thr-close state. The optimal pathway was indicated in blue, whereas suboptimal pathways were depicted in red.

Table S1: Normalized betweenness values connecting the communities in Hsp70-JD system.

| Community #1 | Community #2 | Betweenness value |
|---|---|---|
| 0 | 3 | 1.0 |
| 0 | 5 | 0.543 |
| 0 | 2 | 0.476 |
| 3 | 8 | 0.442 |
| 3 | 6 | 0.431 |
| 0 | 1 | 0.429 |
| 1 | 3 | 0.318 |
| 6 | 10 | 0.290 |
| 2 | 9 | 0.284 |
| 8 | 10 | 0.255 |
| 1 | 8 | 0.232 |
| 7 | 10 | 0.230 |
| 0 | 9 | 0.228 |
| 6 | 7 | 0.227 |
| 4 | 7 | 0.226 |
| 0 | 6 | 0.221 |
| 2 | 7 | 0.217 |
| Cutoff=0.2 | | |
